# Uncovering multiscale structure in the variability of larval zebrafish navigation

**DOI:** 10.1101/2024.05.16.594521

**Authors:** Gautam Sridhar, Massimo Vergassola, João C. Marques, Michael B. Orger, Antonio Carlos Costa, Claire Wyart

## Abstract

Animals chain movements into long-lived motor strategies, resulting in variability that ultimately reflects the interplay between internal states and environmental cues. To reveal structure in such variability, we build models that bridges across time scales that enable a quantitative comparison of behavioral phenotypes among individuals. Applied to larval zebrafish exposed to diverse sensory cues, we uncover a hierarchy of long-lived motor strategies, dominated by changes in orientation distinguishing cruising and wandering strategies. Environmental cues induce preferences along these modes at the population level: while fish cruise in the light, they wander in response to aversive (dark) stimuli or in search for prey. Our method enables us to encode the behavioral dynamics of each individual fish in the transitions among coarse-grained motor strategies. By doing so, we uncover a hierarchical structure to the phenotypic variability that corresponds to exploration-exploitation trade-offs. Within a wide range of sensory cues, a major source of variation among fish is driven by prior and immediate exposure to prey that induces exploitation phenotypes. However, a large degree of variability is unexplained by environmental cues, pointing to hidden states that override the sensory context to induce contrasting exploration-exploitation phenotypes. Altogether, our approach extracts the timescales of motor strategies deployed during navigation, exposing undiscovered structure among individuals and pointing to internal states tuned by prior experience.

## Introduction

Animal behavior emerges from the interplay between external environmental cues and internal states incarnated in complex biological processes such as interoception, neuromodulation and hormonal regulation (Bargmann (2012); Marder (2012)), leading to the emergence of structured behavioral variability across individuals in a population. One major challenge lies in quantitatively assessing how external and internal influences shape this behavioral variability. Most attempts to quantify behavior focus on particular spatiotemporal scales of interest (Schwarz et al. (2015); Marques et al. (2018); Helms et al. (2019)). Recent analysis of short timescale behaviors at high resolution has for example shed light on the neuronal circuits underlying sensorimotor integration (Meng et al. (2024); Kano et al. (2023); Darmohray et al. (2019); Böhm et al. (2016); Knafo et al. (2017); Carbo-Tano et al. (2023)). In contrast, the analysis of long timescale behaviors at low spatiotemporal resolution uncovered changing internal states (Ben Arous et al. (2009); Fujiwara et al. (2002); Flavell et al. (2013); Marques et al. (2020)). However, external and internal influences typically manifest themselves in behavior across several timescales. It is therefore essential to develop holistic approaches that can disentangle the multi-dimensional structure of behavior across scales, from posture movements to search strategies.

Recent advances in machine vision have enabled an unprecedented lens onto the multiple scales of behavior across species (Berman (2018)), allowing for high resolution posture measurements in large environments while spanning several timescales – from milliseconds to hours. We aim to bridge across timescales in behavior by constructing predictive models that can capture statistics of long timescale motor strategies from the integration of fine-scale movements. Markov models offer that possibility in principle: they not only have the potential to be accurate predictive machines, but they also offer a lens into the long-lived properties of behavior through the eigenvalues and eigenvectors of the transition matrix. However, building accurate Markov models requires predictive representations of behavior. Numerous studies in genetic model organisms relied on unsupervised approaches to identify a small number of stereotyped movements, such as “bout types” (Marques et al. (2018); Mearns et al. (2020); Johnson et al. (2020)) or “syllables” (Wiltschko et al. (2015, 2020); Weinreb et al. (2023)). Such behavioral categorization might however unavoidably erase fine-scale information, making the prediction of long timescale sequences challenging. Accordingly, a Markov model built from bout types is unable to uncover the frequency with which larval zebrafish engage in long sequences of bouts (Reddy et al. (2022)), pointing to non-Markovianity. This is not an isolated observation: non-Markovianity appears in behavioral data across species (Schwarz et al. (2015); Berman et al. (2016)) and cannot be decoupled from the behavioral representation.

We thus carefully conceive our analysis to retain fine scale kinematics and history dependence. As recent work has shown (Costa et al. (2023b,a)), taking these aspects into account allows the bridging of fine scale posture movements to longer lived motor strategies within the same Markov model. Using this multiscale approach to behavior, we investigated how sensory information and internal states drive behavior across timescales. We took advantage of the power of larval zebrafish, which are small enough to record multiple individuals for a long duration (Orger and de Polavieja (2017)). We apply our approach on previously collected datasets in which 6-7 days post-fertilization (dpf) larval zebrafish are exposed to a large variety of stimuli (Marques et al. (2018) Reddy et al. (2022)). To design models that accurately capture the long-lived properties of larval zebrafish behavior, we study the dynamical evolution of behavior across maximally-predictive bout sequences. We obtain Markov models that are predictive of the behavioral dynamics of each fish across timescales.

Using parsimonious yet predictive descriptions of an individual animal’s behavior across this hierarchy of timescales enables us to develop a novel approach to dissect individual variability and reveal structure in a large population of animals, disentangling how external sensory cues and persistent hidden states drive behavior. Using models encoding individual behaviors, we reconstruct a phenotypic space that captures overall tendencies for different long-lived motor strategies across fish. We find that the structure of this phenotypic space is only partially determined by sensory contexts, pointing to hidden internal variables. Accordingly, fish with different phenotypic biases exhibit differential sensorimotor transformations across multiple timescales. Surprisingly, we find that prior or current exposure to prey has the most profound impact on behavioral variability. Through simulations, we discover that the persistent phenotypic preferences are particularly well suited for either pursuing and capturing prey, engaging in local searches or performing large distance dispersal, reflecting a previously unexplored exploration-exploitation trade-off.

## Results

### Uncovering the multiple scales of larval zebrafish behavior

Larval zebrafish move via sub-second tail oscillations, referred to as “bouts”, which are separated by periods of rest lasting typically ≈ 0.5 s (Fig. 1A, see Methods). While the tail beats on timescales of *O*(10^−2^ s) during a bout, stereotyped sequences of bouts last several seconds to minutes (Reddy et al. (2022)). We take advantage of large published datasets (Marques et al. (2018)) in which a total of 463 freely-swimming larval zebrafish were recorded from 30 minutes to 3 hours under a variety of sensory contexts. In what follows, sensory context refers to the immediate stimulus experienced by the animal (e.g., navigation in the light, in the dark, looming stimulus, prey capture, etc.), but also the geometry of the arena. Certain groups of fish also experience a different prior context, which involves being raised with live food from 3 days post fertilization (dpf) (see Table. 1). We utilize the posture of the tail during a bout until for a duration of up to 250 ms = 175 frames after bout initiation, so that each bout is encoded as a 8 tail angles × 175 frames dimensional object. To work on concise low dimensional representation of bouts and filter out noisy bout features, we perform principal component analysis (PCA) on all bouts, finding 20 dimensions explain over 95% of the variance across bouts (Fig. S1, see Methods).

**Figure 1.**
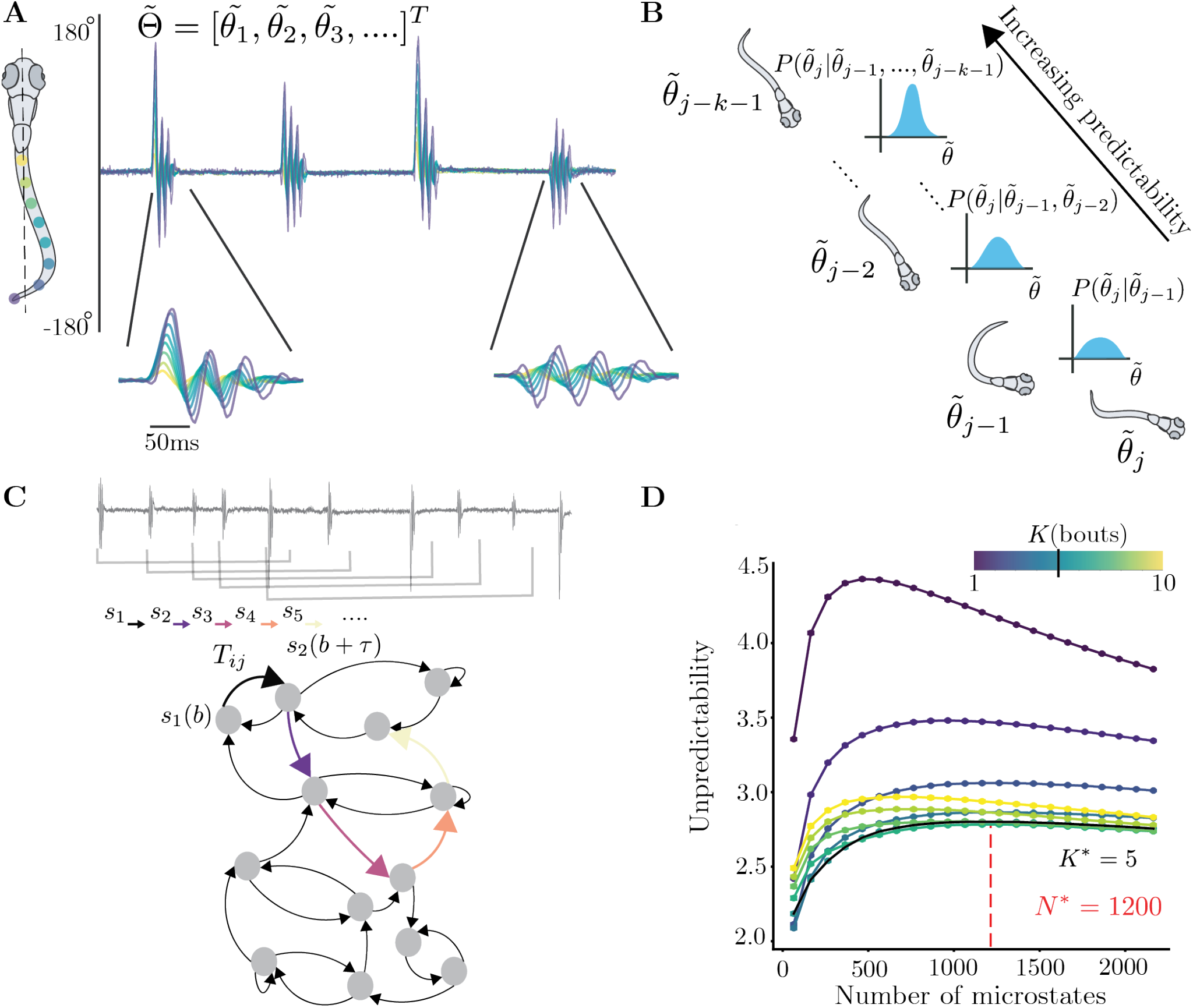
Building a maximally predictive state space for larval zebrafish. **(A)** Typical example of larval zebrafish locomotion bouts, measured as the angle made by points on the tail with respect to the midline. (**below**) Examples of a turn and forward bout. **(B)** In order to recover the underlying dynamics in the bout space, we concatenate bouts from the past behavior of the fish. With more information about the past bouts, the predictability of the future bout is expected to increase. **(C)** The fish’s locomotion is then characterized by a sequence of overlapping windows of *K* bouts (Here *K* = 5). The dynamics of this sequence can be encoded in a Markov chain which we build by discretizing the space of *K* bouts via clustering. The unpredictability of the sequence can then be quantified by the short-time entropy rate of the Markov chain. **(D)** The entropy rates (in nats/bout) of Markov chains for the data from Marques et al. (2018). We sample 7500 bouts from each sensory context equally and estimate the entropy rate of ensemble markov chains at various *K, N*. Note the decreasing entropy rate with respect *K*, minimizing at *K*\* = 5. To maximize the amount of fine-scale dynamics captured by our Markov chain, we pick the maximum number of partitions *N* according to the entropy rate (Shown here at *N* * = 1200 for data from Marques et al. (2018)).

To reveal the long-lived properties of larval zebrafish navigation, we adapt the approach of maximally predictive Markov models introduced in Costa et al. (2023b). We resolve the history-dependence of the dynamics by including past bouts into an expanded representation (Fig. 1B) of the behavioral dynamics. We sample 7,500 bouts from each of the 14 sensory contexts and search for a maximally-predictive representation of the *ensemble* dynamics. We built sequences of bouts (Fig. 1C), increasing the sequence length to maximize the predictability of a Markov chain built from clustering the bout sequences into *N* microstates (see Methods). We assess predictability by estimating the entropy rate of the resulting Markov chain, which reflects the variability in future states given current states as a function of the number of bouts in a sequence, *K*, and the number of microstates *N* (Fig. 1D). Past bouts help narrow down our predictions of the future, and a large enough *N* allows us to retain fine-scale information about each bout’s kinematics. For the dataset from Marques et al. (2018), we set *K*\* = 5 bouts to minimize the entropy rate of the Markov chain, while simultaneously maximizing information content with *N* * = 1200 microstates (see Methods). Given this choice of *K*\* and *N*\*, we then obtain an ensemble transition matrix *T*_ensemble_(*τ*) that captures the overall behavior of all fish across sensory contexts and can therefore reveal long-lived structure in the behavioral dynamics.

The non-trivial eigenvalues *λ*_*k*_ of the ensemble transition matrix *T*_ensemble_ and the respective eigenvectors *ϕ*_*k*_ provide a lens onto the organization of behavior across timescales^1^ (Costa et al. (2023a)). To isolate the long-lived modes from the faster bout-level dynamics, we chose the transition time as *τ* * = 3 bouts (Fig. S2A), obtaining long-lived modes that are well separated from the bulk spectrum (Fig. 2A). As the fish moves, each sequence of bouts corresponds to a value along *ϕ*_*k*_. The time evolution of *ϕ*_*k*_ therefore offers a parsimonious description of the transitions among coarse-grained, longlived behaviors.

**Figure 2.**
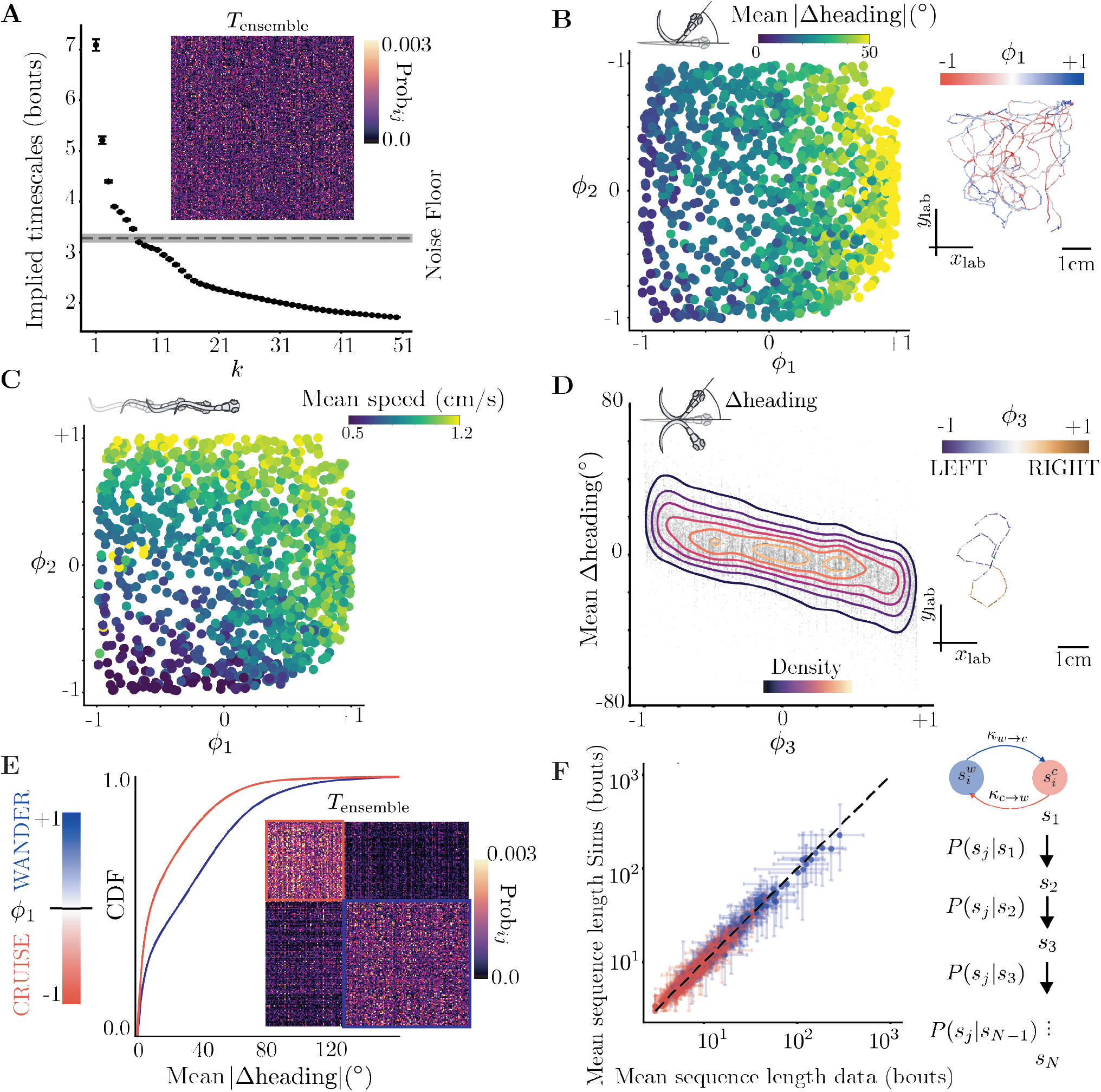
Navigation is driven by 3 long lived modes in a hierarchy of timescales, prioritizing rate of change of heading, speed and egocentric direction bias. **(A)** Implied timescales of the *k*-th mode of *T*_ensemble_, estimated as 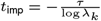, where *λ*_*k*_ are the eigenvalues of *T*_ensemble_ (see Methods). *T*_ensemble_ is built by sampling 7500 bouts from each sensory context (14 total) in Marques et al. (2018) over a 100 seeds. We show the ensemble *T*_ensemble_ as an inset. Error bars represent bootstrapped 95% confidence intervals of the implied timescales over these 100 seeds. **(B)** Microstates organized along *ϕ*_1_ −*ϕ*_2_, color-coded by the mean absolute change in heading. The longest lived mode *ϕ*_1_ seems to correlate to absolute change in heading across bout sequences. **(inset)** An example trajectory of 500 bouts in the lab space color coded by *ϕ*_1_. **(C)** Microstates organized along *ϕ*_1_ − *ϕ*_2_, color-coded by the mean speed. The second longest lived mode *ϕ*_2_ correlates the mean speed across bout sequences. **(D)** The third mode *ϕ*_3_ encodes an egocentric preference for left or right directions. **(inset)** An example trajectory of 40 bouts color coded by *ϕ*_3_. **(E)** These modes can also be used to find an effective coarse-graining of the state space. Partitioning the state space along *ϕ*_1_ reveals *cruising* and *wandering* metastable motor strategies with either low or high changes in heading. Note the block diagonal structure of *T*_ensemble_ organized along the coarse-graining (inset). **(F)** Mean average sequence length of each fish in the data vs simulations. Fish generate highly variable sequence lengths in each strategy, from a few bouts to a few hundreds. Our Markov model built from coarse-grained cruising-wandering states for each fish accurately predicts the mean sequence length in the metastable motor strategies.

We interpret the 3 longest-lived modes by comparing them with common kinematic variables of speed and changes in orientation. The longest-lived mode *ϕ*_1_ roughly correlates with absolute changes in heading direction (Fig. 2B), the second mode *ϕ*_2_ correlates with reorientation and speed (Fig. 2C), and the third mode *ϕ*_3_ with an egocentric direction bias (Fig. 2D) that persists across bouts as previously observed (Dunn et al. (2016)). Note that specifying a single *ϕ*_*k*_ is not sufficient to determine the bout kinematics and the mapping of *ϕ*_1_, *ϕ*_2_ onto specific kinematic variables is non-linear. For example, at intermediate values of *ϕ*_2_, higher values of *ϕ*_1_ may correspond to increased speeds (Fig. 2C). The long-lived modes of larval zebrafish navigation are therefore complex functions of simple kinematic parameters, reflecting joint modes of control on different timescales.

Proceeding from the longest timescales down, we compress the representation of the long-lived dynamics by sequentially identifying *q* motor strategies using an increasing number of faster-decaying eigenvectors of the “ensemble” Markov model across all fish (see Methods) (Fig S2). At the longest timescales, we split along *ϕ*_1_ to identify dynamically coherent behaviors (Costa et al. (2023b)), uncovering 2 novel motor strategies, *“cruising”* and *“wandering”*, corresponding respectively to a low and high rate of reorientation (Fig 2E). The organization of *T*_ensemble_ according to the cruising-wandering categorization reveals a block-diagonal structure indicating the metastability of these strategies(Fig 2E inset). For *q* = 4 motor strategies, our method uncovers *slow* and *fast* variations of *cruising* and *wandering*. For *q* = 7, we obtain left/right variations of slow wandering, fast wandering and fast cruising as well as slow cruising without a left-right bias (Fig S2 D,E,F). To verify that our approach is stable across labs and tracking algorithms, we apply it to a smaller dataset of 6-7 dpf fish exposed to chemical gradients (Reddy et al. (2022)) and tracked with a different software (Mirat et al. (2013)). Our analysis yields similar modes and motor strategies of long-lived behavior (Fig S4, see Methods) indicating that larval zebrafish navigation is organized along a hierarchy of timescales prioritizing rate of reorientation and instantaneous speed.

These metastable states are important motor strategies deployed by the fish across sensory contexts: even though these strategies are obtained from an “ensemble” model, a *q*-state Markov model built for single fish offers a good generative model of its behavior (see Fig. 2F,S3A,B). For a coarse-graining into *q* motor strategies, we simulate the coarse-grained behavior of each fish using its individual transition matrix 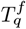 (for cruising-wandering a *q* = 2-state Markov Chain, see Methods). Simulated bout sequences of motor strategies from such transition matrices 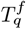 accurately predicted the characteristic sequence length (in bouts) of each strategy for individual fish (for *q* = 2, see Fig 2F and for *q* = 4, 7 see Fig S3A,B). The mean sequence lengths in individual fish vary from a few bouts to hundreds of bouts (Fig. 2F). Such wide range of variation in individual fish is also reflected in the distribution of times spent in either “cruising” or “wandering” across all fish: although cruising persisted 4.04 (4.02, 4.07) bouts and wandering 4.12 (4.09, 4.16) bouts (mean with 95% confidence intervals), these distributions are heavy-tailed (Fig S2G1). There is not one characteristic bout sequence length common across fish and sensory contexts for these motor strategies. Similarly, the bout sequence length of finer scale strategies also varies widely (see Figs S2(G2,G3)). Our analysis yields a hierarchy of Markov models built from an increasing number of coarse-grained states *q* that are *predictive* on increasingly faster timescales (Fig. S3A,B).

### Dissecting the role of motor strategies across sensory contexts

We hypothesize that the large variability among fish could be explained by the usage of these motor strategies in different sensory contexts. As a first assessment, we estimate the probability of visiting different microstates along *ϕ*_1_−*ϕ*_2_ (Fig. 3A, see Methods). While there is variability within each sensory context, fish exhibit preferences for particular motor strategies depending on the sensory context. Fish freely exploring an arena in the light (5 × 5 cm^2^ arena, *n* = 10 fish) mostly perform *fast cruising*: cruising lasts for 5.84 (3.27, 8.78) s and wandering for 1.18 (1.00, 1.93) s (Fig 3A1, median values with 95% confidence intervals). When the geometry of the arena restricts behavior along one axis (1 × 5 cm^2^, *n* = 12), fish still deploy fast cruising for 2.21 (1.97, 2.33) s but also engage in *wandering* for 1.66 (1.35, 2.31) s (Fig 3A2), due to the arena geometry forcing reorientation at the corners (Fig. S6B). In contrast, in response to aversive stimuli, fish mostly performed *wandering* behaviors: fish in the dark (*n* = 37 fish, ≈ 30 minutes) display a preference for *fast wandering* (lasting 8.37 (6.84, 11.90) s) over cruising (lasting 3.95 (2.90, 5.82) s) (Fig 3A3). Similarly, when exposed to an aversive acidic pH gradient localized in space (Reddy et al. (2022)), the avoidance response consists of wandering behaviors (for 2.43 (0.08, 10.89) s, Fig S4E,F) over cruising of 1.99 (0.08, 9.52) s.

**Figure 3.**
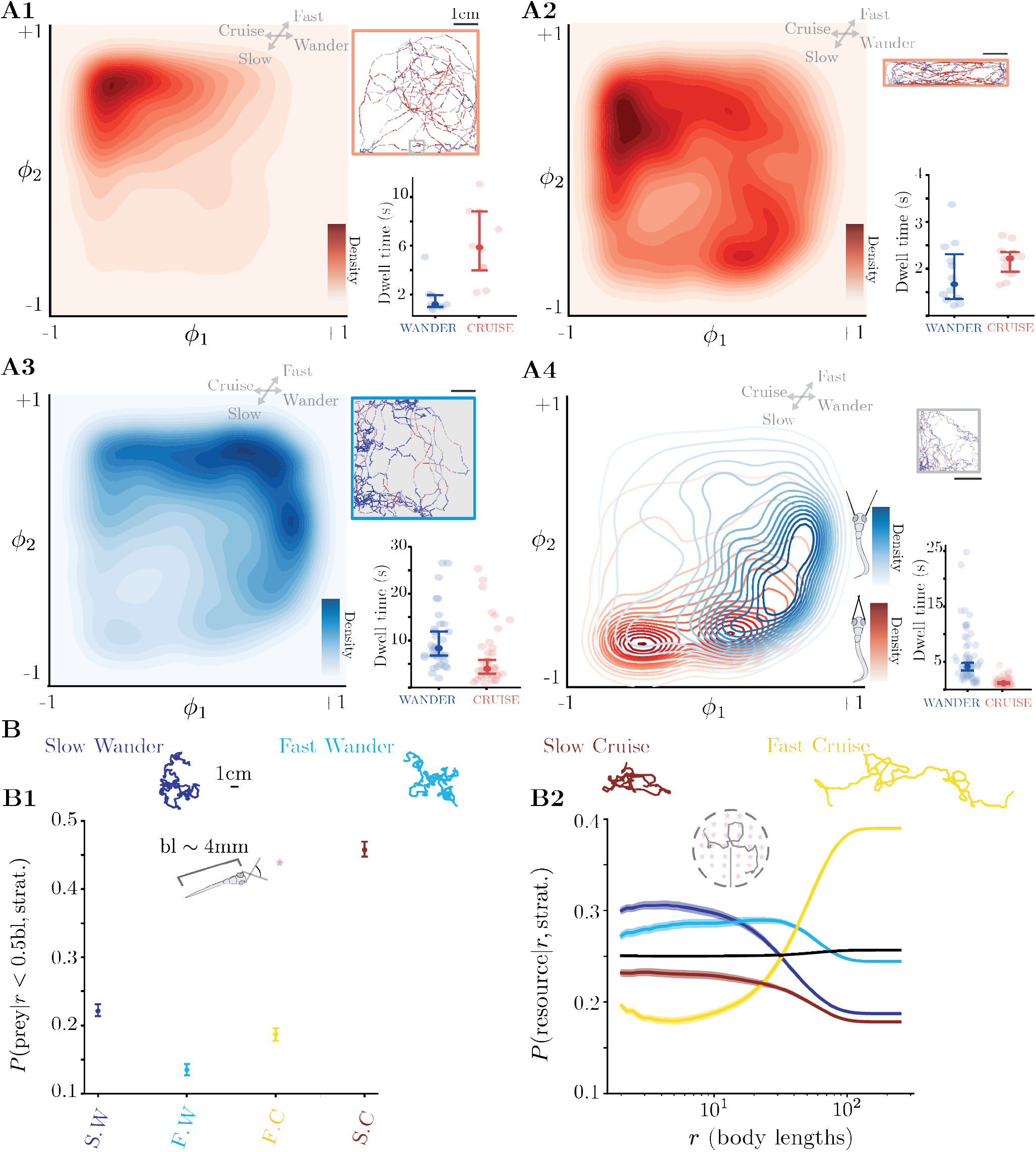
Functional role of motor strategies across sensory contexts. **(A)** Probability density of visiting different bout sequences along *ϕ*_1_ − *ϕ*_2_ (left). As insets, an example fish trajectory color coded by the cruising and wandering states (top right) and the median dwell time in each motor strategy (bottom right), where the error bars represent 95% confidence intervals bootstrapped across the fish belonging to a sensory context (plotted individually in the background). **(A1)** In light (1000 lm/m^2^, 5 × 5 cm^2^ arena, *n* = 10 fish), fish mostly perform fast cruising strategies. **(A2)** In a smaller arena (1000 lm/m^2^, 1 × 5 cm^2^ arena,*n* = 12 fish), we observe a higher usage of wandering strategies, which is particularly enhanced at the short ends where fish are force to quickly reorient (the width of the arena is only about twice the body length of the fish, Fig. S6(B)). This results in a reduction of the time spend cruising when compared with the 5 × 5 cm^2^ arenas in panel A1. **(A3)** Fish in the dark (5 × 5 cm^2^, *n* = 37 fish) show a shift towards fast wandering behaviors. We combine data from two different conditions: the “Dark” condition, in which fish are simply freely-swimming in the dark, and also the first 30 minutes of the “Dark Transitions” condition, in which fish are also freely-swimming in the dark before being exposed to light of different intensities (see Table. 1). **(A4)** In a prey capture assay with ≈ 50 paramecia in the arena (1000 lm/m^2^, 2.5 × 2.5 cm^2^ arena, *n* = 65 fish), fish tend to engage in slow cruising behaviors during eye convergence events, while in the inter-hunt period they mostly perform wandering behaviors. Notably, there is a significant shift from free exploration in the light (A1), with the near absence of fast cruising behaviors even in the inter-hunt period. **(b)** Probability of gathering resources in two distinct regimes: one at short length scales in which the fish has its eyes converged and tries to capture prey within its field of view (60° aperture), and another at large length scales in which we assume the fish can sense resources all around its body (see Methods). We compare the *q* = 4 motor strategies, which correspond to splitting the Cruising and Wandering strategies into their slow and fast variants (Fig S2D). We simulate the dynamics in each motor strategies by inferring transition matrices using only microstates belonging to each of the motor strategies and data from all fish. We then simulate trajectories with a length of 1000 bouts and repeat the process 5000 times starting from random initial conditions. (**B1**) At the scale of less than half the body length, in which the fish begins prey pursuit with its eyes converged (McElligott and O’Malley (2005)), slow cruising is the most successful strategy at acquiring prey. (**B2**) At length scales of larger than twice the body length and while searching all around the body, wandering strategies are effective for short to meso-length scale searching, while fast cruising is effective at large scale dispersal. Black line represents the behavior of an average fish, which is equally efficient at all length scales.

Next, we investigated the motor strategies deployed by larval zebrafish during prey capture. Detailed studies on larval zebrafish hunting behavior have revealed stereotyped sequences comprising of eye convergence promote binocular vision of prey (Bianco et al. (2011); Mearns et al. (2020)) followed by a combination of J-turn, Approach Swim and Capture Swim bout types to successfully track and eat prey (McElligott and O’Malley (2005); Marques et al. (2020); Mearns et al. (2020); Johnson et al. (2020)). In hunting assays, fish noticeably change their navigation: both during prey capture that is estimated by eye convergence but unexpectedly also in between. To capture preys, larval zebrafish hunting paramecia (2.5 × 2.5 cm^2^ arena, *n* = 65 fish) opted for slow cruising (Fig 3A4) that, as expected, primarily comprises J-turns and Approach Swims (Fig S2E). In the inter-hunt exploratory periods, previous analysis relying on bout types suggested similarity with free swimming behavior (Routine Turns and Slow 1/Slow 2 bout types,Marques et al. (2018, 2020); Mearns et al. (2020) or simply exploratory bout types, Johnson et al. (2020)). We uncover that the inter-hunt exploratory navigation strikingly differs from freely swimming associated with fast cruising in the light and is instead dominated by slow wandering (Fig 3A4) (wandering lasts 4.13 (3.11, 4.78) s and cruising 1.14 (1.03, 1.28) s). Our approach indicates that the wandering motor strategy can be deployed for searching in various contexts: either when exposed to aversive cues (darkness: Fig 3A3, acidic pH: Fig. S4) or to appetite cues (prey capture: Fig 3A4).

To dissect what fish can achieve by deploying different motor strategies, we leverage the predictive power of our model to generate synthetic lab space trajectories and assess how distinct motor strategies lead to exploration of space (Fig 3B). We restrict the dynamics to a given metastable strategy and generate artificial 1000 bout-long sequences. We then sample velocity vectors corresponding to these sequences to generate lab space trajectories without boundary conditions (see Methods). Given the simulated trajectories for each motor strategy, we examine two different tasks: one in which the fish has a nearby target within its field of view with the eyes converged, and one in which the fish is broadly searching for resources uniformly scattered on different spatial scales. As expected, slow cruising is the most efficient strategy for pursuing and catching prey with eyes converged (Fig. 3B1). For undirected searches(Fig. 3B2), slow and fast wandering strategies are most efficient on mesoscopic scales, with slow wandering being most efficient up to ≈ 10 body lengths and fast wandering becoming most efficient between ≈ 20 and ≈ 50 body lengths. For resources that are scattered on long distances (beyond ≈ 50 body lengths), fast cruising becomes the most efficient strategy (Fig. 3B2). Our simulation results thus confirm the benefits of distinct motor strategies: fish freely exploring in the light and never exposed to prey mostly engage in fast cruising as a form of long distance dispersal or exploration, while fish pursuing prey engage in slow cruising, and perform slow and fast wandering as effective search strategies or exploitation, either to search for prey, or to escape aversive environments towards safe areas.

### Classifying fish using their behavioral dynamics hints at structure beyond the sensory context

Despite the broad changes in long-lived behaviors induced by the sensory context, we observe large differences between fish for a given context illustrated by the average bout sequence length for each fish in various sensory contexts (Fig. 3A-insets bottom right). Going beyond the *reductionist approach* (Jacobs and Ryu (2023) of focusing on population averages can reveal the structure to the variability among individual, pointing the combined impacts of sensory context and hidden states on behavior. To delineate the effect of sensory context, we first evaluate whether it is sufficient to explain the behavior of every fish by comparing them to each other. This is a non-trivial task as it is not obvious which parameters to select to compare individual animals. Previous studies have utilized heuristic parameters such as handedness, clumpiness, anxiety or boldness/shyness, to measure difference between animals (Werkhoven et al. (2021); Sih et al. (2004). We utilize instead the predictive Markov models of individual fish at different coarse-graining scales to quantify how their behavior unravels over time.

As a direct assessment of the right coarse-graining scale to compare fish, we test whether we can discriminate different sensory contexts based on the behavior of each fish. Intuitively, using too coarse a representation might mix together fish that nonetheless use faster timescale behaviors differently, while using too fine a representation will render each fish unique, thereby challenging the classification task. We classify each fish into their respective sensory contexts by building transition matrices for each fish 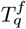 with an increasing number of states *q*, and perform a regularized logistic regression with an 80%-20% train-test split (Fig 4A) (see Methods). The logistic regression shows minimal improvement in accuracy beyond the coarse-graining scale of *q* = 7 metastable strategies that split the fast and slow variations of the *cruising-wandering* strategies into egocentric direction preferences (Fig 4A)(Fig S2C). This result suggests that beyond modulations of reorientation (*cruising-wandering*) and speed (*slow-fast*), persistent *left-right* asymmetry at the experimental timescale also plays a role in distinguishing individual fish from each other across sensory contexts.

**Figure 4.**
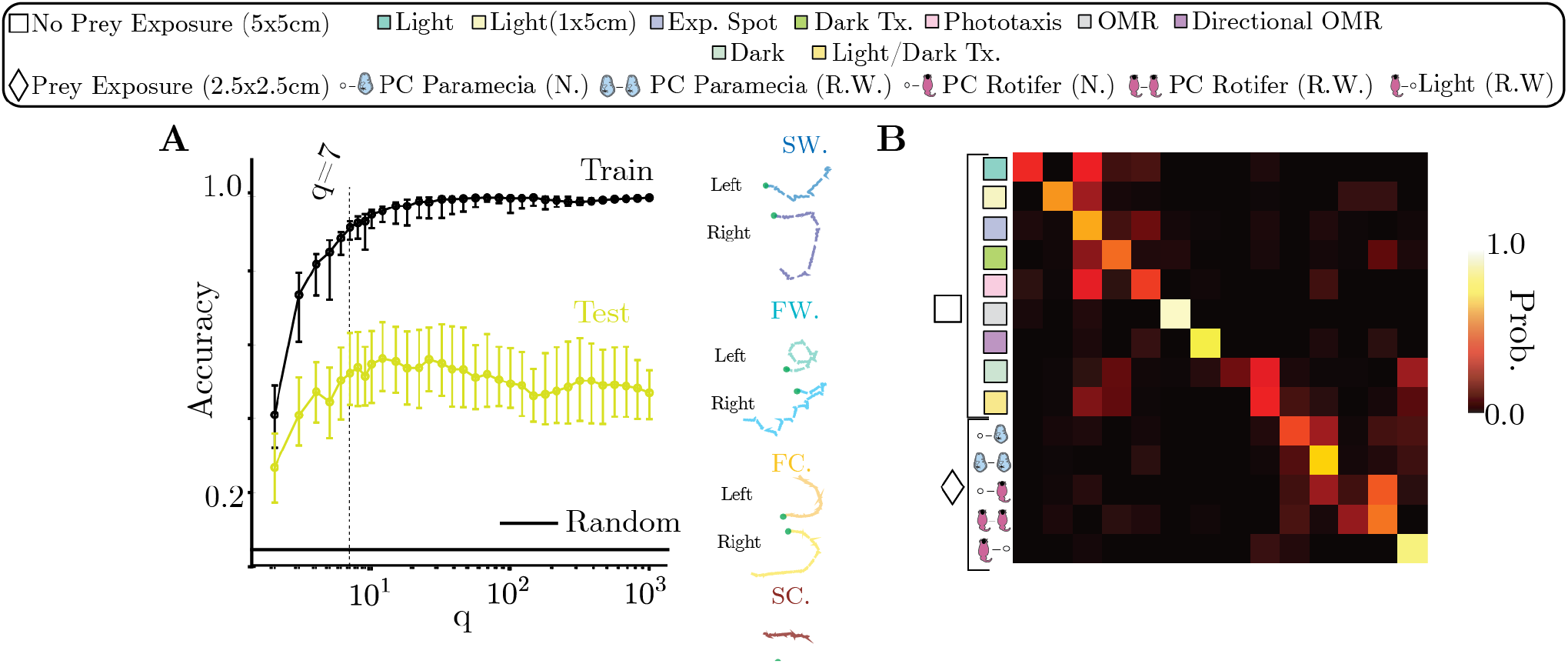
Sensory contexts are insufficient to explain inter-fish behavioral variability. We encode the behavior of each fish in transition matrices 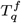 built with an increasing number an coarse-grained states *q*, and use regularized logistic regression to classify each fish into their respective sensory contexts. **(A)** Classification results: accuracy on the Train (black) and Test (yellow) sets as a function of the number of coarse-grained states *q*. We find that the Train accuracy (black) grows continuously as a function of *q* and quickly reaches 1.0. The test accuracy (yellow) is much higher than expected from a random label assignment, but is far from perfect, reaching about 0.5. It starts decaying past *q* ≈ 20 states, but with minimal improvements from *q* = 7 states. This means that while the transition matrix of each fish becomes increasingly unique, allowing for the training accuracy to be nearly perfect, the model quickly fails to generalize to unseen test data: beyond *q* ≳ 7 states the classifier starts to overfit. At *q* = 7 we find Left-Right variations of slow-fast cruising-wandering motor strategies: see example trajectories on the right. **(B)** Confusion matrix of the classifier at *q* = 7: each row measures the proportion of individuals from each context that are classified into every condition. A strong diagonal component indicates that many fish are correctly classified into their respective sensory context, while off-diagonal components also indicate that some fish may get misclassified into a different sensory context, indicating that even in a classifer with a good generalization capacity the behavior of certain fish deviates from the population across sensory contexts.

While the classifier performs much better than random, it is an imperfect predictor of sensory context for a given fish as it only reaches a test accuracy ≈ 50% (Fig 4A). The resulting confusion matrix (Fig 4B), which measures the probability of a fish to be assigned to a given sensory context, has a strong diagonal component, indicating that most fish can be accurately classified – especially for optomotor assays that are wellknown to reliably drive sensorimotor behavior (Severi et al. (2014, 2018)) (center of confusion matrix, Fig 4B). In contrast, most other sensory contexts lead to numerous misclassifications, with some fish behaving more similarly to fish that belong to other sensory contexts.

### Revealing phenotypic groups from inter-fish variability

Our results point to hidden structure in the inter-individual variability that we now reveal with an unsupervised approach. We directly estimate the difference between fish through a modified Manhattan distance among the behavioral phenotypes (encoded in the transition matrices 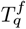 at *q* = 7, see Methods) represented as a distance matrix *D*_*q*_ (Fig 5A). To reveal significant structures in the phenotypic variability, we introduce a novel top-down clustering approach called Hierarchical multiplicative diffusive (HMD) clustering. The finite length of the recordings imposes an effective uncertainty on the measured transition matrices, as two distinct behavioral sequences may result from a finite sampling of the same underlying transition matrix 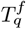. We estimate an effective significance scale for the behavioral phenotype of each fish 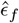 by leveraging simulations of symbolic sequences from 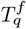, which captures the inherent uncertainty over the precise location of each fish in the phenotypic space due to the finite size of the recordings (see Methods for details). Fig. 5B represents a low-dimensional projection of the phenotypic space obtained using Constant Shift Embedding (CSE, Roth et al. (2003), see Methods): each gray point corresponds to a fish, and two example *q* = 7 transition matrices are depicted in distinct positions of the space (red spots) with the significance scale 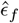 in blue, showcasing how neighboring fish may indeed be behave indistinguishably from each other within the finite size of the recording. Given this effective scale separation among fish, we search for an energy barrier on a family of multiplicative diffusion processes (see Methods). We perform this operation in an iterative fashion, perform a soft clustering along the highest effective energy barrier. This provides a probability of each fish belonging to a given phenotypic group 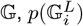 at each iteration *L*. We stop the top-down subdivisioning when the gain in scale separation obtained after subdividing the space becomes negligible (Fig S5B). In Fig. 5C, we illustrate the clustering outcome as a tree diagram, and colorcode each fish in the transition matrix space according to their phenotypic group 𝔾_*i*_,*i* ∈ {1,…, 7}.

**Figure 5.**
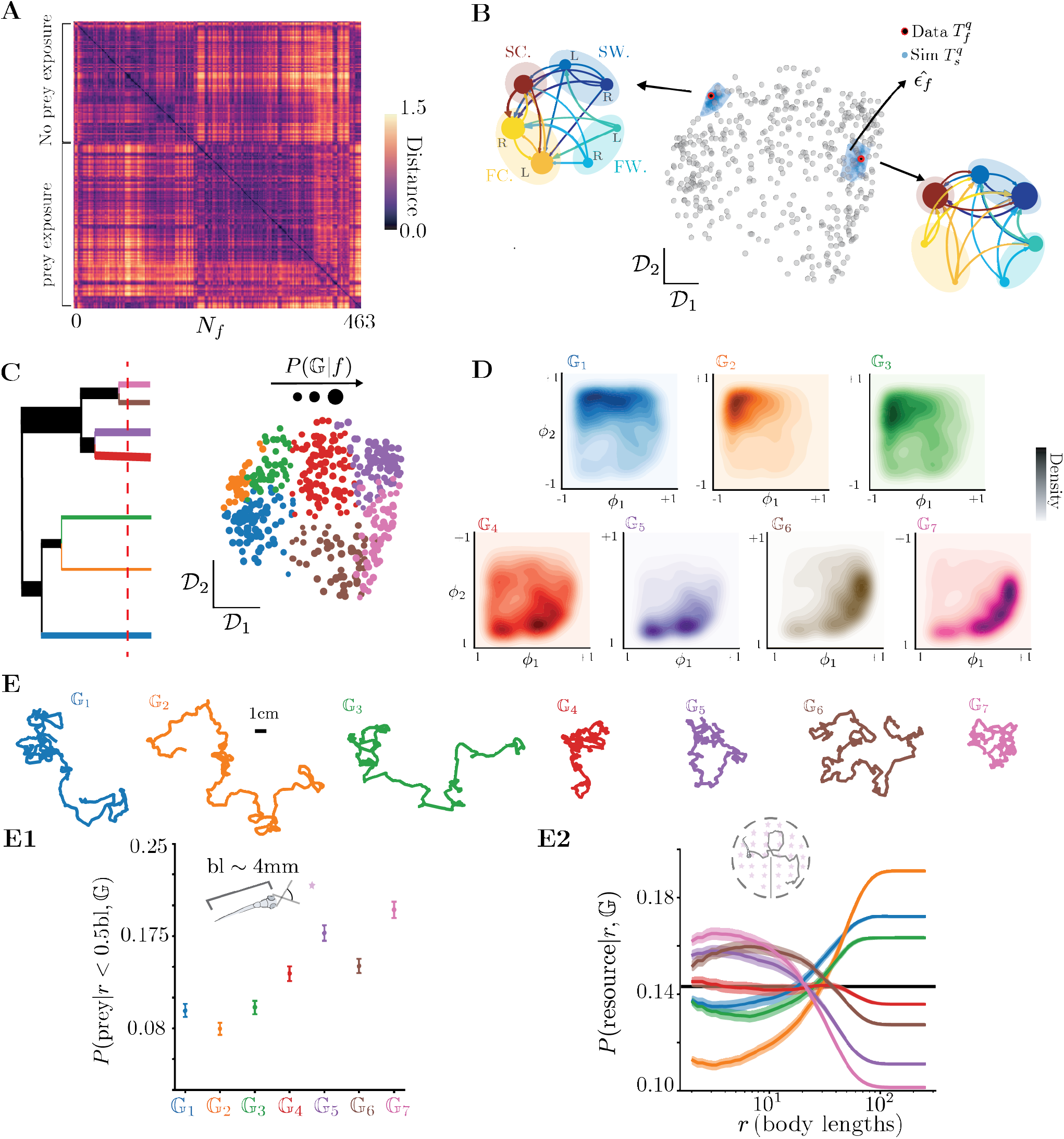
Revealing phenotypic groups by clustering fish based on their behavioral dynamics at the experimental timescale. **(A)** Pairwise distance matrix among the transition matrices of each fish 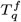 at *q* = 7 states (see Methods). We compare individual fish to each other based purely on their behavioral dynamics (see Methods), finding similaries among fish at several scales. The broad block diagonal structure of the matrix indicates that sub-groups of fish behave similarly, with prey exposure seeming as the broadest determinant of this variability. However, the distance matrix is also highly heterogeneous, suggesting structure beyond the sensory context. **(B)** We use Constant Shift Embedding (CSE) Roth et al. (2003) to embed the phenotypic space into an Euclidean space that preserves the pairwise distances among fish (see Methods). We visualize each individual fish (gray) along the first two dimensions of this *transition matrix space*. We also provide two example transition matrices of fish from different parts of the space. To reveal the multiscale structure to the space of behavioral phenotypes, we estimate whether pairs of fish behave significantly different from each other within the finite time of the experiment (see Methods). We re-estimate transition matrices from finite simulations of each fish (with the same length as the original recordings), and estimate the distance between re-estimated transition matrices and the original transition matrix (blue points surrounding the example fish). Averaging these distances sets an effective scale *‘* within which neighboring fish are indistinguishable from each other (see Methods). **(C-left)** We reveal the structure in the phenotypic space through a top-down fuzzy subdivision of a multiplicative diffusion process, in which distances are rescaled by the effective uncertainty of each fish (see Methods). At each iteration, we identify the group of fish that is most separable (see Methods), and subdivide it. In this way, the ordering of sub-divisions is indicative of the relative scale separation among fish from different groups. We illustrate our clustering approach as a tree diagram. We stop the clustering after 6 levels (7 clusters), the point beyond which the effective distances between fish in clusters stops growing (see Fig S5B). The widths of the branches of the tree are proportional to the number of fish in each cluster. **(C-right)** We color-code each individual fish by their most likely phenotypic group 𝔾_*i*_, where *i* ∈ {1,…, 7}. The size of the dot indicates the posterior probability *P* (𝔾|*f*). (**D**) The clustering reveals significantly different phenotypic groups 𝔾_*i*_, which nonetheless share common behavioral structures according to the hierarchical subdivision. Groups 𝔾_1_, 𝔾_2_ and 𝔾_3_ all exhibit a bias towards fast cruising, but vary in how much they also use fast wandering (𝔾_1_) and slow cruising and wandering (𝔾_3_) strategies. Groups 𝔾_4_, 𝔾_5_ share a bias towards slow cruising and wandering behaviors, with 𝔾_4_ exhibiting a higher proportion of faster cruising. Finally, groups 𝔾_6_, 𝔾_7_ are both biased towards fast wandering strategies, with 𝔾_7_ exhibiting more slow cruising behaviors. **(E)** Probability of gathering resources in two distinct regimes, as in Fig. 3(B) (see Methods). We compare the 7 phenotypic groups, simulating the dynamics in each group by inferring transition matrices across all fish that belong to a particular phenotypic group. We simulate trajectories with a length of 1000 bouts and repeat the process 5000 times starting from random initial conditions. **(E1)** Probability of capturing resources uniformly distributed in a distance shorter than half a body length and within a cone of 60_°_ ahead of the fish (see Methods). Groups 𝔾_5_ and 𝔾_7_ are the most effective at pursuing and capturing prey, reflecting their higher biases towards slow cruising and wandering strategies. **(E2)** Probability of finding resources uniformly distributed in a distance shorter than *r* (see Methods). Groups 𝔾_5,6,7_ are effective for short to meso-length scale searching, while 𝔾_1_ and 𝔾_2_ are most effective at large scale dispersal. Group 𝔾_4_ is approximately equally efficient across a broad range of length scales. Black line represents all groups being equally efficient.

The phenotypic groups reflect the preferences for different motor strategies, Fig. 5D. The main behavioral difference emerging from the top-down clustering is the use of fast cruising for 𝔾_1,2,3_. Conversely, fish in groups 𝔾_4,5,6,7_ opt for slow cruising and wandering behaviors. Further iterations reveal finer scale preferences for motor strategies. In Fig. S5C we also report mean dwell times of fish conditioned on the group 𝔾 they belong to. To further assess the role of preference in motor strategies, we evaluate how they lead to a differential exploration of space. We proceed as in Fig. 3(B), and simulate artificial fish trajectories in each behavioral group using group level transition matrices *T*^*g*^ (see Methods). We find that groups 𝔾_7_ and 𝔾_5_ are most efficient at pursuing nearby prey (Fig. 5E1). Along with 𝔾_6_, these groups are also efficient at searching for resources uniformly scattered on distances up to 10 body lengths (Fig. 5E2). In contrast, group 𝔾_2_ is optimized for long distance dispersal, becoming most efficient on distances beyond ≈ 30 body lengths. Notably, we also find that group 𝔾_1,3,4_ display average efficiency for searching at short length scales. At longer length scales 𝔾_1,3_ become more efficient, while 𝔾_4_ drops. We thus discover that the structure of the phenotypic space reflects an exploration-exploitation trade-off taking place at the level of the population.

### Phenotypic groups capture behavioral differences imposed by sensory context

We now assess how much of the structure in the phenotypic space can be explained by sensory context. We place the average phenotype of each sensory context on the phenotypic space (Fig. 6A1), and estimate how many fish in a given sensory context belong to each behavioral group in Fig. 6A2. We find that sensory contexts can drive preferences for different regions of the phenotypic space, but no single sensory context completely maps onto a single phenotypic group. To elucidate this further, we evaluate how sensory contexts split at different levels of the hierarchical clustering process. Fig. 6B shows the probability of belonging to a given group 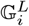 at different subdivision levels *L*. The first split *L* = 1, which captures the main axis of variation, mainly differentiates naive fish from fish exposed to prey (Fig 6B1). Consequently, the largest phenotypic variation favoring high exploitation emerges upon exposure to prey, changing the behavior at the timescale of experiment to promote wandering and slow cruising behaviors. At *L* = 4 subdivisions Fig 6B2, which splits 𝔾_4,5_ from 𝔾_6,7_, broadly separates fish raised with rotifers but freely swimming in the light (Fig 6B2 below in blue) away from fish in hunting assays (Fig 6B2 below in red). The prior hunting experience impacts even the freely swimming behavior of the fish, resulting in a phenotype that is different not just from naive fish but also from fish in hunting assays. At *L* = 5 subdivisions Fig 6B3, we mostly split fish hunting for paramecia versus rotifers. Notably when naive fish are placed in hunting assays for either paramecia or rotifers, the behavioral phenotypes of the fish spread across all groups. However, when fish are raised with paramecia from 3 dpf on, they are typically biased towards higher exploitation 𝔾_5_, in contrast with that fish raised with rotifers which mostly belong to 𝔾_4_. Altogether, our approach uncovers how different prior exposure to prey leads to different emergent hunting phenotypes at a later developmental stage.

**Figure 6.**
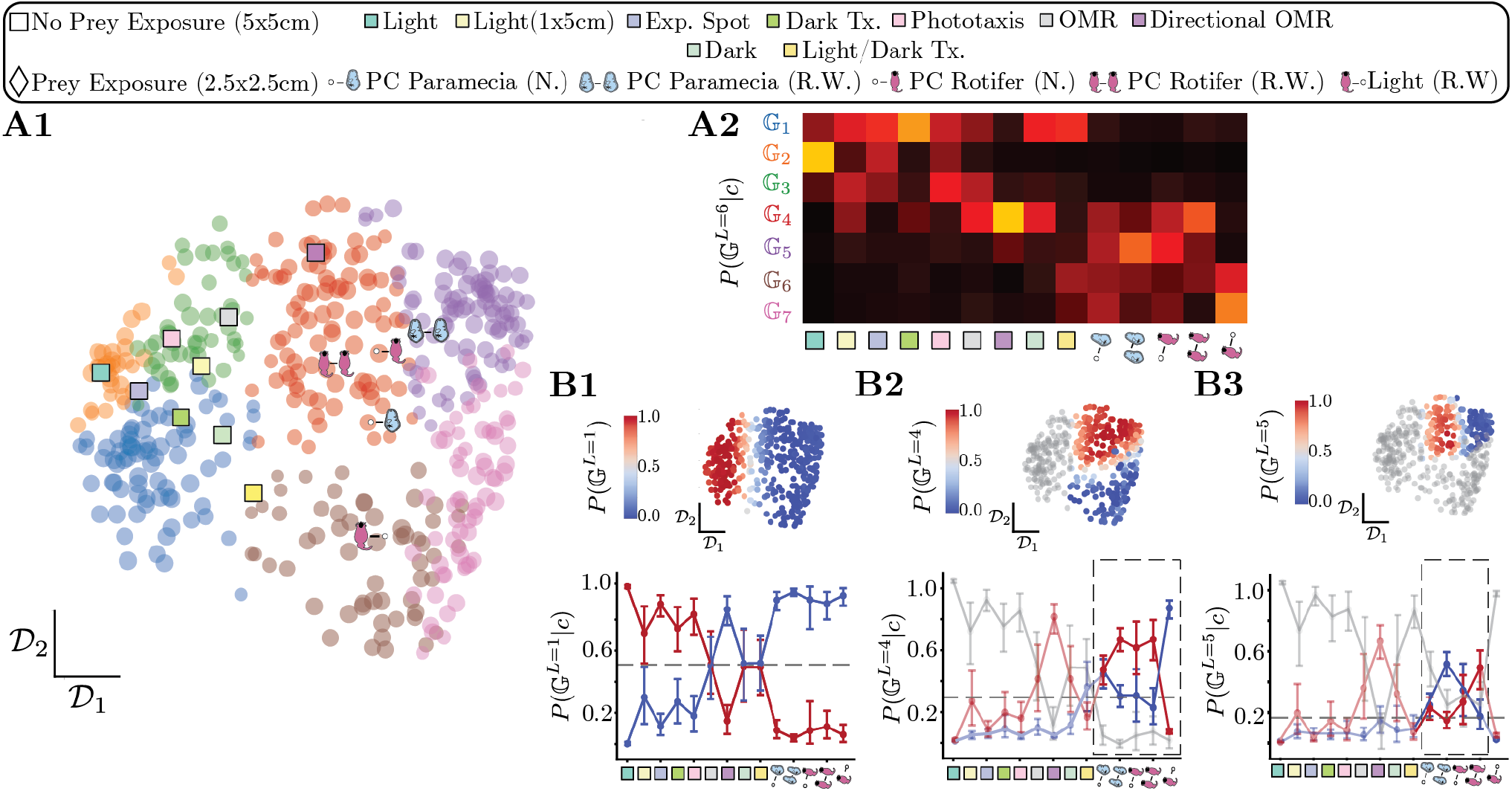
Phenotypic group structure explained by the sensory context, revealing prey exposure as a major determinant of variation. **(A1)** Transition matrix space color-coded by the 7 phenotypic groups. We also show the position of the average transition matrix in each behavioral assay, using the same markers as in Fig. 4. **(A2)** Probability of belonging to group 𝔾 for fish in a given experimental condition *c, P* (𝔾^*L*=6^|*c*). Note that while no single sensory context maps onto a phenotypic group completely, the majority fish in certain contexts have preferences for certain groups. For example, most fish never exposed to prey belong to 𝔾_1,2,3_, while fish exposed to prey belong to groups 𝔾_4,5,6,7_ **(B)** To delineate this further, we show the probability of belonging to group 𝔾 for fish in a given experimental condition *c* at different iterations in the hierarchical subdivision *L*. **(B1)** The first split *L* = 1 neatly distinguishes fish that were exposed to prey (either during or prior to the assay) from fish that were never exposed to prey. Notice also that the directional OMR condition is closer to the prey capture conditions, indicating that the sensory contexts in this assay somewhat emulate the spontaneous behavior in the prey capture assays. The OMR, Dark and Light/Dark transitions assays are distributed across the two main groups. **(B2)** At the 4th iteration, *L* = 4 fish that were previously exposed to prey but are in freely-swimming conditions neatly split from the fish that were in a hunting assay. **(B3)** At the 5th iteration, *L* = 5 fish in hunting assays that were raised with different types of prey mostly belong to different groups. Interestingly, naive fish that are hunting for the first time split equally, whereas fish that were raised with either paramecia or rotifers mostly belong to distinct groups.

To further assess the differences in behavior among naive and rotifer-raised fish, Fig 6B2, while taking into account the effects of the arena geometry, we quantify the spatial distributions of different motor strategies. We find that in freely-exploring conditions, naive fish wander mostly only at the corners of the arena where they are forced to reorient (Fig S6A,B) – both in the 5 cm × 5 cm and in the 1 cm × 5 cm arenas. In contrast, fish raised with prey and freely-swimming in a 2.5 cm × 2.5 cm arena performed wandering throughout the entire arena (Fig S6C,D) and were more frequently oriented towards the wall, suggesting that wandering is local search strategy to leave the arena rather an an anxiety driven wall-hugging behavior (Fero et al. (2011); Schnörr et al. (2012)).

### Phenotypic groups reveal persistent hidden states impacting behavior across timescales

Sensory contexts captures some of the variation in the phenotypic space, but we also find a large amount of variability within sensory contexts (Fig. S7), pointing to hidden variables that impact the sensorimotor transformations performed by the fish. We therefore investigate whether such variability can reflect inner motivational states by investigating the variability observed upon exposure to prey. We focus on fish hunting for paramecia (bold highlight, Fig. 7). Half the fish in this assay do not belong to the population average groups 𝔾_4_ and 𝔾_5_. As a hunting indicator, we leverage the measurement of time spent with eye converged throughout the experiment (Bianco et al. (2011)). Fish belonging to exploration phenotypes (𝔾_1,3_) rarely hunt, whereas fish whose phenotype is tuned to pursuing prey (𝔾_5,7_) hunt often (Fig 7A2). Interestingly, fish raised with rotifers but freely swimming without prey in the light (Fig 7B1) also show differences in their eye convergence rates that depend on the phenotypic groups: fish tuned to exploration (𝔾_1_) barely attempt hunts, whereas fish that are more tuned to exploit (𝔾_7_) exhibited a higher rate a eye convergence (Fig 7B2). Overall, while our approach does not directly take into account the prey capture rate, the structure to the variability reveals evidence for motivational states that drive contrasting exploration-exploitation phenotypes.

**Figure 7.**
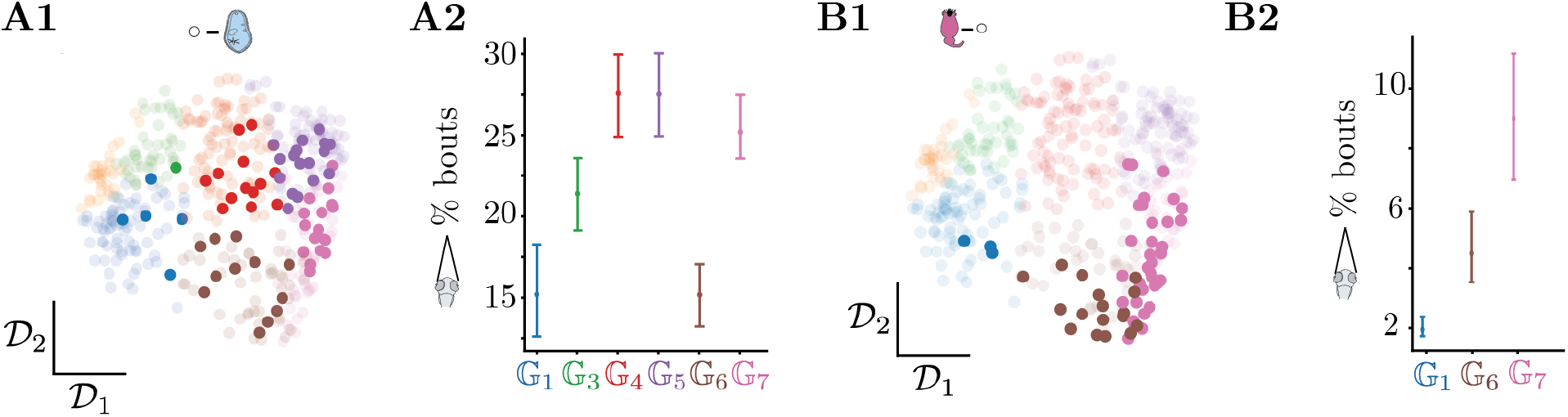
Phenotypic group structure reveals differences in sensorimotor transformations among fish within a sensory context. **(A1)** Transition matrix space color-coded by the 7 phenotypic groups, of naive fish hunting for paramecia (represented in bold). Note the high degree of variability in the dynamics of the fish within this condition. **(A2)** Percentage time spent in eye convergence for naive fish hunting paramecia, structured by the phenotypic group 𝔾. Note that fish in 𝔾_1_ and 𝔾_6_ have lower time spent with eyes converged, indicating that these fish are potentially hunting lesser than fish in other groups. **(B1)** Transition matrix space color-coded by the 7 phenotypic groups, of fish freely exploring in the light, but raised with rotifers (represented in bold). **(B2)** Percentage time spent in eye convergence across fish in the light raised with condition, structured by the phenotypic group 𝔾. Note that while the overall time spent in eye convergence is quite low as these fish are freely swimming, fish in 𝔾_7_ still have significantly higher time in eye convergence, pointing to differing hidden states impacting behavior within this condition.

## Discussion

In this study, we aim to reveal how behavioral phenotypes arise as a function of sensory inputs and latent variables that impact behavior on multiple timescales. Via the analysis of maximally-predictive bout sequences in larval zebrafish, we reveal a hierarchy of three long-lived modes of navigation organized by timescale: a long lasting mode that reflects the rate of reorientation, a faster mode that also encodes for speed and a third mode that captures egocentric direction preferences. We then discover phenotypic preferences along these modes among individuals which seem to be partially explained by the sensory context (arena size, illumination and experimental stimuli). To quantify the structure and origin of this variation, we compare individual fish to reveal phenotypic groups using only the behavior at the experimental timescale. Through *in silico* experiments we find that the phenotypic group structure corresponds to exploration v/s exploitation trade-offs apparent across individuals. These groups reveal that a major driver of phenotypic variation is the exposure to prey, engaging a strong preference for local exploitation. This variation not only impacts the dynamics deployed by animals in hunting assays but also affects the freely swimming behavior of fish in the light. However, the structure of the phenotypic space also reveals similarities between fish in distinct sensory contexts, pointing to a combined impact of sensory context and persistent hidden variables that differ among individuals. We show that phenotypic group structure conditions the sensorimotor transformation at short timescales by promoting either exploitation or exploration, showcasing how these latent variables prevail over the immediate sensory contexts.

### A hierarchical organization of timescales in larval zebrafish behavior

Sensory-evoked navigation consists in chaining locomotor bouts in response to external sensory cues from the environment and internal states. We reveal long-lived modes of behavior by constructing Markov models from bout sequences deployed by the larval zebrafish, accounting for the history dependence in the behavior and the maximal possible variability in posture dynamics of bouts (Costa et al. (2023b)). Our method consistently provided a hierarchy of motor strategies, reflecting rate of reorientation, speed and directionality, in distinct datasets acquired by different users, from different arenas, and laboratories. Changes in long-lived modes corroborate observations from previous studies noticing changes in reorientation rates in the dark (Horstick et al. (2017)), speed modulation during optomotor response (Severi et al. (2014)) or recurrence in left or right bias during navigation in the light (Dunn et al. (2016)). However, these modulations were previously observed by focusing on a specific kinematic parameters, choosing a specific time-window to analyze, or classifying all bouts into a limited repertoire (such as left versus right for (Dunn et al. (2016)). Our unbiased analysis expands this view by quantifying the dynamics of behavior as the chaining rule of maximally predictive bout sequences, thereby providing a complete and simultaneous picture of the multiple axes of behavior and revealing how they are organized hierarchically by timescale, which could have been missed upon focusing on specific kinematic parameters.

Methodologically, we use the inferred long-lived modes of a transfer operator to effectively identify an increasing number of motor strategies that capture shorter lived behaviors (Fig. 2). Coarse-graining behavior using the non-trivial eigenvectors of transfer operators is equivalent to using the Information Bottleneck approach (Schmitt et al. (2023)), commonly applied in behavioral analysis (Berman et al. (2016)). This equivalence holds for behavioral representations that evolve in a Markovian fashion, stressing the importance of working with maximally predictive bout sequences (Costa et al. (2023b,a)). Additionally, it is common to define discrete states by clustering similar movements, identifying movements that are executed often, or by building Hidden Markov Models (HMM) (Marques et al. (2018); Johnson et al. (2020); Mearns et al. (2020); Wiltschko et al. (2015, 2020); Calhoun et al. (2019)). However, the level of discreteness required is challenging to define and interpret with such approaches. To avoid this issue, we use the notion of timescale separation as a guiding principle for defining when a given discrete representation holds, which we reveal by coarse-graining according to the *dynamics*. On the longest timescales, cruising-wandering strategies provide a good-enough representation. To capture faster timescale behaviors, we utilize a increasing number of finer scale motor strategies. From these motor strategies, we build transition matrices for individual fish that are predictive of each fish’s behavior across coarse-graining scales. This parsimonious encoding of the behavioral program of each fish quantifies its behavioral phenotype, providing us with specific timescales in the motor strategies deployed by individuals. If these motor strategies were arbitrarily defined, simple Markov models would not make good predictions.

The predictive power of our approach is adequate but has some limitations: while we can predict whether the next bout sequence will belong to a coarse-grained motor strategy, we cannot predict the precise kinematics of each bout in the sequence. This may simply reflect an inherent bound to predictability coming from the stochastic nature of the behavioral dynamics, or be due to insufficient data. Further work will resolve this question. While our approach is effective at capturing behavioral dynamics across several timescales, there are opportunities for enhancement: i) longer behavioral recordings in larger arenas with novel sensory contexts will likely reveal new long-lived behaviors; ii) we could include the spatiotemporal properties of the stimuli and the arena geometry, which impacts the dispersal properties of the fish’s behavior (Fig S8); iii) to capture any slow non-stationary changes to behavior, we could introduce time-varying transition rates as was done in Costa et al. (2024); iv) we could incorporate the inter-bout intervals in our approach as we find them correlated with the bout sequences used (Fig S2 h1-h3), similar to a previous observation (Johnson et al. (2020)).

### Sensory contexts drive overall preferences for motor strategies

Behavior in larval zebrafish is classically studied to investigate instantaneous sensorimotor transformations. By quantifying the behavioral responses of a large population of fish in different sensory contexts, our approach uncovered biases of motor strategies across multiple timescales in response to diverse sensory contexts. We show that larval zebrafish display a preference for fast cruising when freely swimming in the light, while in aversive settings they display preferences for wandering. Previous work discovered that upon exposure to darkness, larval zebrafish spiral for 2-3 minutes as a local search and turn more in the dark (Horstick et al. (2017)). Our results confirm these observations, revealing long-lived wandering states in the dark (≈ 8 s) interspersed by short segments of cruising (≈ 4 s), so that fish mostly wander in dark environments (≈ 30 min), (Fig 3A3). Our simulations indicate that higher wandering corresponds to the fish performing more local area searches. The intermittent switches to cruising could help the fish to disperse to newer areas faster than if the local search strategy dominated the entire behavior, as also previously hypothesized (Horstick et al. (2017)).

Recent studies have identified stereotyped bout types used during hunting sequences through unsupervised clustering (Marques et al. (2018); Johnson et al. (2020); Mearns et al. (2020)), and analysis of fish dispersal within pre-selected time-windows enabled the discovery of long-lived “exploration” and “exploitation” states in hunting assays (Marques et al. (2020)). We find that the hunting sequence is represented in slow cruising, which constrains the variation in cruising timescales (Fig. 3A4). In the inter-hunt period, fish mostly engage in wandering strategies which could correspond to fish actively searching for prey in their surroundings. Thus beyond the tight stimulus-response loop of the hunting sequence, the behavior of the fish throughout the experiment is overall attuned towards searching and hunting. Furthermore, our comparative analysis over multiple sensory contexts differentiates the inter-hunt exploratory period from navigation in the light, during which fish mostly display fast cruising. Fast cruising may thus represent an alternative “exploration” strategy that is extremely rare upon exposure to prey. In *C. elegans*, exploration-exploitation behaviors have been linked to minutes long “roaming” and “dwelling” states respectively (Fujiwara et al. (2002); Flavell et al. (2013)): the “roaming” state is dominated by faster forward locomotion with rare reorientations, while the “dwelling” state is characterized by smaller scale movements that do not coherently engage the whole body resulting in low speeds Fujiwara et al. (2002); Flavell et al. (2013); Hebert et al. (2021). We hypothesize that “slow cruising”, with its slow speeds and targeted reorientations may be analogous to the “dwelling” state of *C. elegans*, while “roaming” recalls “fast cruising” behavior, which is also dominated by forward bouts and sporadic routine turns and is most efficient at long distance dispersal. Our approach offers an opportunity to further dissect the neuromodulatory control mechanisms driving these exploration-exploitation trade-offs by taking into account the finer scale bout sequence dynamics, thus acting as a powerful complement to previous approaches that quantified the overall displacement of the fish within pre-selected time-windows (Marques et al. (2020)).

By providing a simultaneous handle on the multiple timescales that govern behavior such as cruising and wandering, differential modulation of speed and egocentric direction preference, our approach sets the stage for dissecting the underlying brain circuits for navigation in a transparent vertebrate brain. Along with the serotoninergic modulation of motor activity mentioned previously, we can dissect how multiple brain regions function together to give rise to this hierarchy of timescales in behavior. The hindbrain oscillator, also called anterior rhombencephalic turning region has been identified as one brain region that confers a persistence of the left/right steering in the larval zebrafish (Dunn et al. (2016); Wolf et al. (2017)). We hypothesize that the interplay of this region with the mesencephalic locomotor region (MLR, Carbo-Tano et al. (2023)), the nucleus of the medial longitudinal fasciculus (Wang and McLean (2014); Berg et al. (2023)), projecting onto the reticular formation (Orger et al. (2008); Carbo-Tano et al. (2023)) could explain the dynamics we uncover. In particular, the modulation of the MLR by dopamine released from posterior tuberculum (Carbo-Tano et al. (2023)) could explain the long-lived persistence of cruising behaviors. There is already some evidence for this in mice, where dopamine release has been associated with long time scale (seconds to minutes) persistence of motor sequences (Markowitz et al. (2023)).

### Variability in multiscale behaviors exhibits hierarchical structure

While it is apparent that sensory contexts give rise to preferences for motor strategies, it is not clear how these contexts shape the structure of phenotypic variation. This is further confounded by the large amount of individual variability, evidenced in our inability to predict the sensory context of many individuals based on their behavior alone (Fig 4). We hypothesize that the variability in emergent phenotypes is driven by a combination of sensory contexts and persistent hidden states. To study the structure of this variability, we introduce an unbiased approach that compares individual animals directly using their multiscale behavior to reveal phenotypic groups. Recent studies developed strong insight into the structure of behavioral variability by comparing important kinematic parameters (Werkhoven et al. (2021)), the probability of the occurrence of behaviors (Hernández et al. (2021)), or the parameters of minimal behavioral models (Helms et al. (2019); Goc et al. (2021)). Our approach for comparing individuals incorporate all of these aspects: 1) the hierarchy of timescales in the ensemble dynamics naturally reveals the important behavioral parameters in the form of interpretable motor strategies; 2) the predictive power of our coarse-grained Markov models allows us to encode the dynamics of the behavioral transitions for each individual, capturing also the probability of visiting each behavior; 3) our ability to scan across coarse-graining scales reveals the precise amount of fine-scale information needed to compare animals.

Quantifying structure in individuality is a non-trivial task due to the uncertainty associated with the variable duration and fish swimming frequency across recordings. This uncertainty renders bottom-up agglomerative clustering approaches unreliable. We solved this issue by turning this limitation into a feature, introducing a novel top-down clustering algorithm that directly leverages the uncertainty in the estimate of each behavioral phenotype to provide an effective scale separation between individuals. We obtain a soft cluster assignment, which directly highlights the interplay between a continuous and discrete phenotypic groups. While this clustering method was developed for analyzing behavior in freely moving animals, it could be generalized to the study of other dynamic biological processes where variability around common principles is the hallmark, such as cell migration (Brückner et al. (2020)) or neural dynamics (Brynildsen et al. (2023)).

### Behavioral phenotypic groups emerge as consequence of both sensory drives and persistent hidden states

Prey capture is an innate behavior as larval zebrafish at 6-7 dpf must feed to ensure their survival (Wilson (2012)). Previous exposure to prey has been shown to increase their rate of capture initiation (Oldfield et al. (2020); Lagogiannis et al. (2020)). The first split in the top-down clustering process is largely driven by exposure to prey (Fig 6A1): most fish exposed to prey perform local exploitation (𝔾_4,5,6,7_) while naive fish perform long distance dispersal (𝔾_1,2,3_). This exposure to prey has a even stronger impact on the behavioral phenotype of fish than aversion to darkness (Fig S7). Remarkably, even fish previously raised with prey but recorded freely exploring in the light belong to exploitation groups (𝔾_6,7_). Such long lasting preference may be due to recruitment of the hypothalamus during hunting, initiating a feeding state in the fish(Muto et al. (2017)). Serotoninergic (Filosa et al. (2016)) and dopaminergic signalling (Zaupa et al. (2024)) may also be implicated in extending duration of this feeding state, manifesting as a large phenotypic variation in behavior. We also find that the nature of the preys (paramecia or rotifers, Marques et al. (2018))) leads to distinct behavioral phenotypes suggesting that fish may adapt their motor strategies to the kinematics of the prey. Our work opens the possibility for future studies investigating the dynamics of predator-prey interactions in greater detail.

Previous studies investigating individual variability in *D. melanogaster* found evidence for persistent hidden states (Hernández et al. (2021); Werkhoven et al. (2021)). Here, we report that the large variability among individuals pointing to hidden states drives variation along these exploration-exploitation type phenotypes.

While the exposure to prey induces exploitation-related phenotypes, we also find a significant number of fish belonging to exploration phenotypes (and vice versa for fish never exposed to prey). One can speculate on what these hidden states may correspond to. Fish in hunting assays that deviate from the population average also display lower rates of eye convergence, possibly due to different motivational states such as feeding or arousal that can profoundly impact sensorimotor integration (Pantoja et al. (2016, 2020)).

Our work provides novel approaches for the quantitative study of phenotypic variability across timescales. We reveal how this variability is structured by the sensory context of the animal and hidden motivational states that drive the animals along an exploration-exploitation trade-off. Our approach lays the groundwork for the study of the sources of individual variability, and can be easily extended to other species. Combined with recent advances in large scale tracking of wild animals, we offer a new avenue to study how phenotypic variation is shaped by the environment, serving the cause of ecology and biodiversity.

## Methods

### Software availability

Code and data for reproducing our results is publicly available at https://github.com/GautamSridhar/Markov_Fish

### Data collection and preprocessing of the data from Marques et al. (2018)

We collected datasets investigating larval zebrafish behavior from Marques et al. (2018). In the dataset from Marques et al. (2018), wild-type Tubingen zebrafish larvae (*Danio rerio*) were used at 6-7 days post fertilization. Larvae were recorded at a high temporal resolution (700Hz), at two different pixel sizes (58*µ*m in the 5 × 5 cm^2^ arenas and 27*µ*m in the 2.5 × 2.5 cm^2^ arenas) and tail-tracking of 8 points on the tail was performed online using custom software (Marques et al. (2018)). From this dataset, we collect 463 fish from 14 varying sensory contexts (Table 1). In Marques et al. (2018), we retained all detected bouts that generated a velocity of at least 4 mm/s (equivalent to one body length of the fish). We also noticed that the detected bout start and bout end cut-off were slightly inaccurate and many detected bouts did not showcase the full motion of the tail from start to end. To account for this, we took 10 extra frames from before the start of the bout as the new bout start frame. For the new bout end, we chose the frame up to when the fish was generating a velocity of at least 4mm/s (equivalent to 1 body length), or 175 frames (equivalent to 250ms), whichever limit came first. We provide further details for each behavioral assay below:

#### Light

**5** × **5** cm^**2**^: Fish were presented with a uniform light from below with an illuminance of 1000 lm/m^2^ in a 5 × 5 cm^2^ squared arena with 3 mm of depth.

#### Light

**1** × **5** cm^**2**^: Fish were presented with a uniform light from below with an illuminance of 1000 lm/m^2^ in a 1 × 5 cm^2^ squared arena with 8 mm of depth.

#### Dark

**5** × **5** cm^**2**^:Fish were presented with darkness (0 lm/m^2^) from below in a 5 × 5 cm^2^ squared arena with 3 mm of depth.

#### Expanding Spot

**5** × **5** cm^**2**^: An expanding dark spot at different speeds (0.25, 0.5, 1, 1.5, 2.0, 2.5 cm/s) and different orientations (0°, 90°, 180°, 270°) were presented in closed loop 4 cm away from the larva. Stimuli were randomized and presented every 2 min. This assay was done in a 5cm x 5cm squared arena with 0.3 cm of depth.

#### Dark Transitions

**5** × **5** cm^**2**^: The spontaneous swimming with light transitions assay was based on Burgess and Granato (2007). Fish were left in the dark for 30 minutes and then presented with uniform light at different intensities (0, 12, 44, 104, 232, 447, 790, 1890, 4700 lm/m^2^) for 3 min in a 5 × 5 cm^2^ arena with 3 mm of depth.

#### Phototaxis

**5** × **5** cm^**2**^: The phototaxis assay was based on Huang et al. (2013) and performed in 5 × 5 cm^2^ arena with 3 mm of depth. The stimulus consisted on a uniform brightness of varying intensity (100, 410, 780, 1250 lm/m^2^) on one side of the fish and darkness on the other. The stimulus was in closed loop with the larva for a duration of 12 s.

#### Forward Optomotor response

**1** × **5** cm^**2**^: The forward optomotor response assay was performed as described in Severi et al. (2014), but using a 1 × 5 cm^2^ with 8 mm depth arena. Drifting gratings with a spatial period of 1 cm and ten different speeds (0, 2.5, 5, 7.5, 10, 15, 20, 30, 40, 50 mm/s) were presented from below when the larva was at the extremities of the arena. Trials would end when fish reached the opposite end of the arena. After, there were 5s of intertrial interval of homogenous light (1000 lm/m^2^) and a new trial would start with gratings in the opposite direction. In trials that larvae were not able to reach the opposite end of the arena (*>* 30 s) a 10 mm/s grating was displayed until it swam the remaining distance.

#### Directional Optomotor response

**5** × **5** cm^**2**^: The directional optomotor response assay was performed as in Orger et al. (2008). A **5** × **5** cm^**2**^ arena with 3 mm of depth was used. Drifting gratings with a spatial period of 1 cm, moving at 10 mm/s, and from 24 different orientations (15° apart) were presented from bellow to the larva and in closed loop with its orientation. Trials lasted 10s and started when the larva was in the center of the arena. During the inter-trial interval, that would last at least 5s, circular converging gratings were projected to drive the larva to center of the arena.

#### High Lux light-dark transitions

**5** × **5** cm^**2**^: Fish were exposed to alternating 3 min periods of high illuminance light (5000 lm/m^2^) and darkness (0 lm/m^2^) light in an 5 × 5 cm^2^ arena with 3 mm of depth.

#### Prey Capture

**2.5** × **2.5** cm^**2**^: All prey capture datasets were performed in 2.5 × 2.5 cm^2^ arenas with 3 mm depth and larvae were illuminated from above (1000 lm/m^2^). Fish were fed with 50-100 paramecia (*Paramecium caudatum*) or rotifers (*Brachionus plicatilis*) and were allowed to hunt for 1-2h. A subset of fish were never fed until the assay, while others were raised, starting at 3 days post fertilization, with the type of prey that was used in the assay. To ensure that these fish were not satiated they were starved for at least 2 hours before the assay.

### Freely exploring in the light, reared with rotifers

**2.5** × **2.5** cm^**2**^: Fish were fed with 50-100 rotifers starting at 3 days post fertilization and then placed in 2.5 × 2.5 cm^2^ arenas with 3 mm depth and illuminated from above (1000 lm/m^2^) to freely swim. To ensure that these fish were not satiated they were starved for at least 2 hours before the assay.

### Data collection and preprocessing of the data from Reddy et al. (2022)

In the dataset from Reddy et al. (2022), wild-type AB zebrafish larvae are recorded at 7 days post fertilization. Larvae were placed in arenas of size 14 cm × 1 cm × 0.4 cm and recorded for 10 minutes at a temporal resolution of 160 Hz and pixel size of 70*µ*m. Half the fish in each experiment were exposed to acidic (pH ≈ 2) gradients at both horizontal ends of the arena. After diffusion, the acidic solution formed a steep gradient 20 mm away from each end of the arena. 8 points on the fish’s tail were tracked using Zebrazoom (https://zebrazoom.org/) and recordings were processed as indicated in Reddy et al. (2022). We retain 218 fish in this dataset which performed more than 250 bouts during the recording. Similar to the previous dataset, we keep all bouts up to either the detected bout ends, at most till 40 frames (250ms).

### Principle component analysis (PCA) on the space of bouts

By measuring 8 tail angles over *N*_frames_ (175 frames for the data from Marques et al. (2018) and 40 frames for the data from Reddy et al. (2022)), we end up with a high dimensional *N*_frames_ × 8 representation of each bout.To increase tractability while reducing noise, we perform a principle component analysis (PCA), Fig. S1. For the dataset of Marques et al. (2018), we randomly sample 25 recordings 180 times (amounting to ≈ 50, 000 bouts per sample) and then calculate the eigenvectors and eigenvalues of the covariance matrix built along the bouts of this recording. Across multiple resamples of the data, we received very stable eigen-values as represented by the miniscule 95% errorbars in Fig S1A. We then average the covariance matrices across these multiple resamples and use the eigenvectors of the averaged covariance matrix as the eigenvectors to represent the feature space of bouts (Fig S1B). We retain 20 dimensions for the dataset from Marques et al. (2018) which covers upwards of 95% of the cumulative variance explained (**Fig S1a**). Similarly, for the dataset from Reddy et al. (2022), we retain 12 principle components.

### Maximally predictive state spaces for larval zebrafish behavior

To study the dynamics in the bout space, we adapt the method described in Costa et al. (2023b,a). Given *l* bouts (where one bout is a *d* dimensional object after PCA) in a recording, we stack *K* bouts together in overlapping windows to receive a matrix of size (*l* − *K* + 1) × *dK*. This process is repeated for multiple values of *K*.

In order to approximate the dynamics in a high dimensional delay-embedding space, we rely on a discrete approximation of the transfer operator as described in Costa et al. (2023b) -we first partition the bout sequence space through *k-means* clustering for which we utilize the scikit-learn function with *k-means ++* initialization (Pedregosa et al. (2011)). After partitioning the space into *N* microstates, the dynamics is recapitulated through a transition matrix *T*_ensemble_ that is built by sampling bouts equally from each sensory context. We then check for which value *K*\* the short time entropy of rate of *T*_ensemble_ minimizes. The entropy rate is measured as *h* = −∑_*ij*_ *π*_*i*_*T*_*ij*_ log *T*_*ij*_, where *π*_*i*_ represents the *i*-th entry of the invariant density, obtained as the first eigen-vector of *T*_*ij*_ (see Costa et al. (2023b) for further details). We then search for *K\** such that *∂*_*K*_*h*(*K*\*)∼0, Fig. 1(d),S4. We similarly select the number of partitions *N* = *N* * for the correct *K* = *K*\* at the maximum point just before finite-size effects cause a reduction the short-time entropy rate (Fig. 1(D),S4). This allows us to incorporate as much information about the fine-scaled dynamics as possible. In this fashion, we have moved from a high-dimensional continuous space of bout evolution to a Markov chain representing the dynamics as a sequence of *N* * symbols.

For the datasets from Marques et al. (2018), we utilize 463 fish from 14 different sensory contexts (**Table 1**). We estimate the entropy rates by building transition matrices *T*_ensemble_, and bootstrapping across a random sampling of 7500 bouts from each condition over 100 random seeds, to ensure a uniform sampling across sensory contexts. For the dataset from Reddy et al. (2022), we have 2 sensory contexts. The entropy rate reflects a bootstrapping across a random sampling of 40,000 bouts from each condition over 50 random seeds. Once *K*\* and *N*\* were chosen, we selected the cluster labels that maximize the entropy rate with respect to the random resampling, and use such cluster labels for the rest of the analysis.

### Building an ensemble transition matrix *T*_ensemble_

To study the dynamics of maximally predictive bout sequences, we estimate an ensemble transition matrix *T*_ensemble_ from the available data. At *K* = *K*\*=5 and *N* = *N*\* = 1200, we randomly sample 7500 bout sequences from each condition over 100 seeds and estimate *T*_*s*_(*τ*) for varying transition times · for each seed *s*. We estimate the implied timescales of the reversibilized transfer operator from the eigenvalues of each re-Versibilized 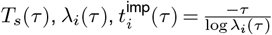, and choose *τ* = *τ*^*\**^ = 3 bouts such that the fine-scaled dynamics have relaxed to the steady state distribution, isolating the long-lived modes (see Costa et al. (2023b) for further details), Fig. S2A. We calculate bootstrapped estimates of the implied timescales across the 100 estimates of *T*_*s*_(*τ*). The noise floor is calculated by shuffling the symbolic sequence of the dynamics, re-estimating reversibilized transition matrices and obtaining their largest real eigenvalue. Finally, we estimate the ensemble *T*_ensemble_ by averaging the transition matrices *T*_*s*_(*τ*\*) and row-normalizing.

### Operator based partitioning into metastable strategies

The eigenvectors of the reversibilized transition matrix *T*_ensemble_ can be used to provide an effective coarse-graining of the dynamics into metastable strategies via the formulation of almost-invariant sets (Froyland and Dellnitz (2003); Froyland (2005); Costa et al. (2023b). Each eigenvector of the reversibilized *T*_ensemble_ provides an ordering of the behavioural state space along a particular timescale. To find the first level of coarse-graining into the two metastable strategies of cruising-wandering, we rely on the fact that the first eigenvector provides an optimal 2-way cut of a graph encoding the dynamics (Froyland and Dellnitz (2003); Froyland (2005)). In practice, we define two macroscopic sets (which correspond to collections of microstates *N*_*i*_) by splitting along *ϕ*_1_,

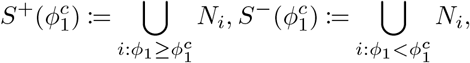

where 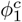 is a threshold that is chosen to maximize the metastability of a set. We measure the metastability of each set *S* by estimating how much of the probability density remains in *S* after a time scale *τ*,

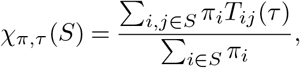

where *π* is the the invariant density, estimated directly from *T*. To estimate the overall measure of metastability across both sets *S*^+^ and *S*^−^, we define

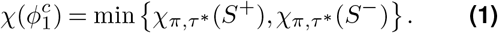

which we maximize with respect to 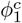. See Costa et al. (2023b) for further details and applications to known dynamical systems. In Fig. S2B we show the overall coherence measure as a function of *ϕ*_1_, which we use to find the 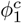 that maximizes coherence, defining “cruising” and “wandering” states.

Subsequent coarse-graining into *q* shorter timescale strategies is achieved by adapting the *q-way* cut formulation from Froyland and Dellnitz (2003), which relies on performing a *k-means* clustering using the first non-trivial ⌈log_2_ *q*⌉ eigenvectors of the reversibilized transition matrix. We introduce a new heuristic for obtaining *q-way* cuts that respect the metastability of the dynamics. We first identify the value of each *ϕ*_*k*_ that maximizes the coherence Eq. 1, defining *S*^+^ and *S*^−^ by thresholding along each eigenvector *ϕ*_*k*_. We then transform each eigenvector by subtracting the 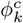 that maximizes the coherence, obtaining a new 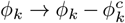 that is centered at 0. We also normalize negative and positive values such that the eigenvector loading becomes equally spaced on the interval [−1, 0] and [0, 1]. All eigenvectors *ϕ*_*k*_ of the reversibilized *T*_ensemble_ in Figs. 2,3,5,S2 are reported after performing these transformations. Finally, we use the notion of kinetic maps Noé and Clementi (2015); Noé et al. (2016) to convert dynamics in the space of these eigenvectors into equivalent distances in Euclidean space using this subsequent transformation,

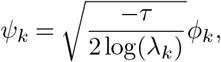

where *λ*_*k*_ is the eigenvalue corresponding to *ϕ*_*k*_and *τ* is the transition time of the dynamics. We identify *q* motor strategies by performing *k-means* clustering to on *k*-dimensional space defined by the transformed eigenvectors *ψ*_*k*_, *k* ∈ [1, ⌈log_2_ *q*⌉], weighted by the steady state invariant measure of each microstate.

The *k-means* clustering is performed using the *scikit-learn* python package Pedregosa et al. (2011) over 1000 automatic repetitions with a *k-means ++* initialization. Coarse-graining at *q* = 2, 4, 7 is reported in Fig. S2. Further coarse-graining for a larger number of states is used in Fig. 4.

### Analyzing the kinematics encoded long-lived modes of *T*_ensemble_

We find correlations between the long-lived eigen-vectors *ϕ*_*k*_ and interpretable kinematic variables in Fig. 2(B,C,D). For this, we calculate the mean of each kinematic variable in the stacks of *K* = *K*\* bouts that belong to a particular cluster *s*_*i*_, *i* ∈ {1,…, *N* *}. Similarly for the inter-bout interval in Fig. S2(H), we report the median value of the distribution of inter-bout intervals belonging to a particular cluster. For the coarse-grained strategies, we collect every trajectory of different fish executing one of the metastable strategies and bootstrap 100 times to calculate the cumulative distributions functions of the mean absolute change in heading and the mean speed with error bars in Figs. 2(E),S2(D). For the preference for egocentric direction in Fig. S2(F), we assign each bout in a trajectory as left or right if the change in orientation from the start of one bout to the next is positive or negative respectively. The ratio of left or right movements in a trajectory is then calculated, and bootstrapped 100 times to report the mean value with error bars.

### Simulations of symbolic sequences

We use individual transition matrices of each fish at various coarse-graining scales 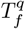 to simulate the symbolic dynamics of the fish. This translates to learning 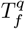 for each fish from its individual symbolic sequence at a particular *q* and then evolving the symbolic dynamics according to 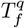 from its start state. We repeat this process a 100 times for each fish. We then calculate the mean sequence length in these simulations in each strategy at *q* = 2, 4, 7 by bootstrapping across the 100 simulations for each fish. (Fig. 2(F),S3).

### Sampling lab space trajectories from symbolic sequences

In order to simulate artificial trajectories from symbolic sequences, we sample velocity vectors 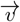 (of the head position of the fish from the start of one bout to the next) from the distribution of vectors 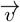 of a particular microstate *N* = *N* *. Thus for the symbolic sequences of real data, we sample 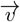 according to the distribution of velocities within each discrete state *s*_*i*_(*t*) for increasing times *t* (Fig S8A). To compare simulations with data in Fig. S7, we utilize transition matrices *T* with all the microstates *N* = *N* * at *τ* = 1 instead of *τ* = 3, which provides access to the fine-scale dynamics while retaining the information about the long-lived modes. We calculate this transition matrix *T* using the ensemble of fish imaged in a 5 × 5 cm^2^ arena in order to minimize mixing boundary effects from velocity vectors in different arenas. Finally, we simulate 1000 bout long symbolic sequences according to this transition matrix *T*, sampled from random initial conditions, and sample velocity vectors 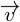 from the underlying distributions within each if the samples microstates. Three example trajectories are shown in Fig S8B)

For the mean squared displacement (MSD) in Fig. S8C, we report the MSD estimated from real trajectories (Real data in black from fish is in a 5 × 5 cm^2^ arena), artificial trajectories that effectively remove boundary conditions by sampling velocity vectors from the real symbolic sequence of the ensemble of fish (Real seq. in lime) and artificial trajectories obtained from simulated symbolic sequences (Sim seq. in magenta). In addition, we report the result of enforcing reflective boundary conditions on simulated symbolic sequences, limiting the arena size to 5 × 5 cm^2^ (Refl. Sims in red).

For the long simulations from metastable strategies at the coarse-graining scale *q* = 4, we estimate an ensemble transition matrix restricted to the microstates that belong to the metastable strategy, using *τ* = 1, and sample from the inferred transition matrix 1000-bout long symbolic sequences 5000 times. Using the obtained symbolic sequences, we then sample from the underlying velocity vector distributions to get artificial lab space trajectories. The mean squared displacement calculations are reported in Fig S8D. For simulations from behavioral groups 𝔾 = *g*, we similarly build ensemble transition matrices *T*^*g*^ at *τ* = 1 of fish belonging only to a specific group and then simulate 1000-bout long symbolic sequences 5000 times and sample velocity vectors as before.

### Examining the preference for metastable strategies across conditions

To investigate the preference for a strategy in a particular sensory context, we project the symbolic sequence of all fish in that condition along *ϕ*_1_, *ϕ*_2_. We then calculate a histogram of this projection with 50 bins along each axis, smoothed by a Gaussian kernel with a window size of 3 bins.

### A distance metric to compare transition matrices

Given a fish’s individual transition matrix 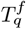 at a particular coarse graining scale *q*, we aim to compare dynamics of remaining in a particular metastable state or transitioning out of it, with that of another fish with transition matrix 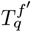. We use the *L*1 norm (Manhattan distance) between every row of the transition matrices 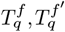 to calculate these distances. With a distance value for each row of the two matrices 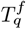 and 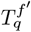, we then calculate an average of these distance values to receive one value to compare fish to each other. Formally speaking, we have

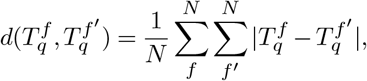

which qualifies the criteria for being a metric.

### Constant Shift Embedding for distance matrices

Using the distance metric defined above, we are able to compare distances across 463 fish and build a distance matrix *D*_*q*_ at a particular coarse-graining scale *q* (datasets from Marques et al. (2018)). We embed the calculated distance matrix between fish into a space where the Euclidean metric preserves the distances between fish using a Constant Shift Embedding (CSE)(Roth et al. (2003)). Unlike methods like Multidimensional Scaling (MDS) (Kruskal (1964)) which minimize a cost function, CSE relies on solving an eigen-vector problem, where each eigenvector for eigenvalues greater than 0 correspond to the dimensions of the Euclidean embedding. The eigenvalues of the CSE operation provide us with importance weights for each eigen-vector.

### Logistic Regression to determine the correct coarse-graining scale to compare fish

We attempt to classify each fish into its respective sensory context using only its behavior. We do so by comparing the test accuracy of a logistic regression at different levels of coarse-graining *q*, from 2 metastable strategies to a 1000. To do this, we first build individual transition matrices 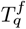 of all fish *f* at a particular coarse-graining scale *q* and transition time *τ*. The coarse-graining *q* is discovered in the same way as previously described from the eigenvectors of the ensemble transition matrix *T*_ensemble_. We then estimate a distance matrix *D*_*q*_ among all 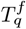, and perform CSE to embed it in a Euclidean space. We project the distance matrix to all the eigenvectors discovered by CSE with eigenvalues greater than 0.0. We then train logistic regressions (using the scikit-learn package Pedregosa et al. (2011)) over 100 shuffles of the data using a 80%-20% train-test split. Since each sensory context in the dataset from Marques et al. (2018) contains different number of recordings, we shuffle recordings within each sensory context for every random seed and then sample 80% of fish from each context and leave 20% for testing. The logistic regression loss is weighted to account for this class imbalance. For each run of the logistic regression, we perform a grid search for the correct value of the *L*2 regularization parameter *–* using a 5-fold cross-validation (all of these steps are performed automatically by the scikit-learn GridSearchCV module). The model with highest validation accuracy is subsequently applied on the test data. We then compare the mean accuracy over the training and test sets (all weighted by the sample weight) over 100 shuffles of the data (Fig 4A). We also extract confusion matrices for each run of the logistic regression over a shuffle of the data and report the mean confusion matrix across the seeds (Fig 4B).

### Hierchical multiplicative diffusive (HMD) clustering

After selecting the correct coarse-graining scale *q* = 7 based on the logistic regression, we work with the distance matrix *D*_*q*_ obtained with *q* = 7. We introduce a novel approach to perform a top-down clustering of the transition matrix space that takes into account the uncertainty in the estimate of the distances between data points. We do this by estimating an effective significance scale for each fish 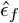 This amounts to reestimating transition matrices 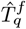 from a 100 different simulations of symbolic sequences from 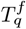 and calculating the average distances between the 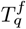 and the re-estimated 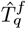 Thus we have

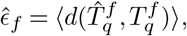

In Fig S5A, we provide the 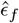 for each fish. For a single fish *f*. With this effective significance scale, we effectively rescale the distance among fish, and treat the top-down clustering as a search for metstability in a multiplicative diffusion process defined by the kernel

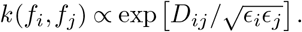

This operator rescales the distances between data points based on their uncertainty, such that points that have overlapping significance scales are effectively closer to each other than data points that do not overlap. For the first iteration of the hierarchical clustering, we leverage the first non-trivial eigenvector of this operator (the Fiedler vector) that partitions the data points into two (Froyland and Dellnitz (2003)) by finding the location of an effective barrier height in the diffusion dynamics. This is effectively equivalent to performing spectral clustering into two clusters using a 1-D diffusion map of data. To obtain a fuzzy cluster assignment, we discover this clustering by fuzzy c-means clustering (*skfuzzy* package Warner et al. (2019)). We repeat this process iteratively: At the second iteration, we first create two multiplicative diffusion operators using only the data in each cluster from the first iteration. We then quantify the effective barrier height of these diffusive processes by measuring the level of metastability of the diffusive dynamics within the cluster. Effectively, this is calculated in the same way as the dynamics, by maximizing the coherence along the first non-trivial eigen-vector of the diffusion operator. Then we split the cluster that maximizes the metastability. Because we discover a fuzzy clustering assignment, we essentially receive posterior distributions for each iteration. However, since these posteriors are limited the cluster that was split, we extend them other data points by estimating distances between the split cluster center and the unseen data points and converting them to probabilites. This is handled automatically by the skfuzzy package.

### Estimating kinematic parameters for phenotypic groups 𝔾

We convert the posterior *P* (𝔾 = *g*|*f*) to *P* (*f*|𝔾 = *g*) using Bayes rule and calculate subsequent kinematic parameters as expectation values. For example, for the case of Fig 7, we calculate the percentage time spent in eye convergence *ec* in a sensory context *c* as -

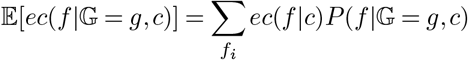

We estimate errorbars by utilizing importance sampling, where we repeatedly estimating the mean of *ec*(*f* |𝔾 = *g, c*) by drawing *f* from *P* (*f* |𝔾 = *g, c*).

## Author Contributions

G.S., M.V., A.C.C and C.W. conceived and designed the project. G.S. and A.C.C performed the analysis. G.S, J.C.M and M.B.O curated and preprocessed the datasets. G.S., A.C.C and C.W. wrote the paper. All authors reviewed the results and approved the final version of the manuscript.

## Acknowledgements

We would like to thank all the members of the Wyart lab for extensive discussions, especially Dr. Martin Carbo-Tano, Dr. Elias T. Lunsford, and Mahalakshmi Dhanasekar for for their insight in fish neuroanatomy and behavior. G.S was supported by the European Union’s Horizon 2020 Research and Innovation program under the Marie Skłodowska-Curie Grant No. 813457 awarded to C.W and managed by Dr. Joana Guedes (https://zenith-etn.com) at the Paris Brain Institute (Institut du Cerveau). G.S, C.W and A.C.C were also supported by Fondation Bettencourt-Schueller (FBS-don-0031), the European Research Council (ERC Consolidator ERC-CoG-101002870), the team grant Fondation pour la Recherche Médicale (FRM-EQU202003010612), the Agence Nationale pour la Recherche (ANR) LOCOCONNECT, RocSMAP, ASCENTS, MOTOMYO. G.S and C.W also acknowledge support in part by the grant NSF PHY-1748958 of the Kavli Institute for Theoretical Physics (KITP) and the Gordon and Betty Moore Foundation Grant No. 2919.02. A.C.C was supported by LabEx ENS-ICFP: ANR-10-LABX-0010/ANR-10-IDEX-0001-02 PSL* and M.V by the NIH Grant No. 1RF1NS128865-01. J.C.M received the support of a fellowship from “la Caixa” Foundation (ID100010434, LCF/BQ/PR21/11840005) and from the Portuguese Foundation for Science and Technology (EXPL/MED-NEU/0957/2021). M.B.O. was supported by grants from the Volkswagen Stiftung “Life?” Initiative (A126151) and ERC (NEUROFISH 773012).

## Footnotes

1 Prior to estimating the eigenvalues and eigenvectors of the transition matrix, we enforce its reversibility so that *T*_ensemble_ refers to as a “reversibilized” transition matrix, whose eigenvectors optimally distinguish metastable states (Dellnitz and Junge (1999); Froyland (2005); Froyland and Padberg (2009); Costa et al. (2023b)).

