## Supplementary Table and Figures for "Uncovering multiscale structure in the variability of larval zebrafish navigation"

### Supplementary Information

**Table 1.** Different Sensory conditions from Marques et al. (2018)

| Sensory Condition | Number of fish | Arena Size (W cm x H cm x D cm) |
|---|---|---|
| Light | 10 | 5 x 5 x 0.3 |
| Light | 12 | 1 x 5 x 0.8 |
| Expanding Spot | 61 | 5 x 5 x 0.3 |
| Dark Transition | 27 | 5 x 5 x 0.3 |
| Phototaxis | 30 | 5 x 5 x 0.3 |
| Forward OMR | 12 | 1 x 5 x 0.8 |
| Directional OMR | 30 | 5 x 5 x 0.3 |
| Dark | 10 | 5 x 5 x 0.3 |
| High Lux Light/Dark Transitions | 22 | 5 x 5 x 0.3 |
| Prey Capture Paramecia Naive | 69 | 2.5 x 2.5 x 0.3 |
| Prey Capture Paramecia Raised With | 98 | 2.5 x 2.5 x 0.3 |
| Prey Capture Rotifer Naive | 15 | 2.5 x 2.5 x 0.3 |
| Prey Capture Rotifer Raised With | 31 | 2.5 x 2.5 x 0.3 |
| Light Rotifer Raised With | 60 | 2.5 x 2.5 x 0.3 |

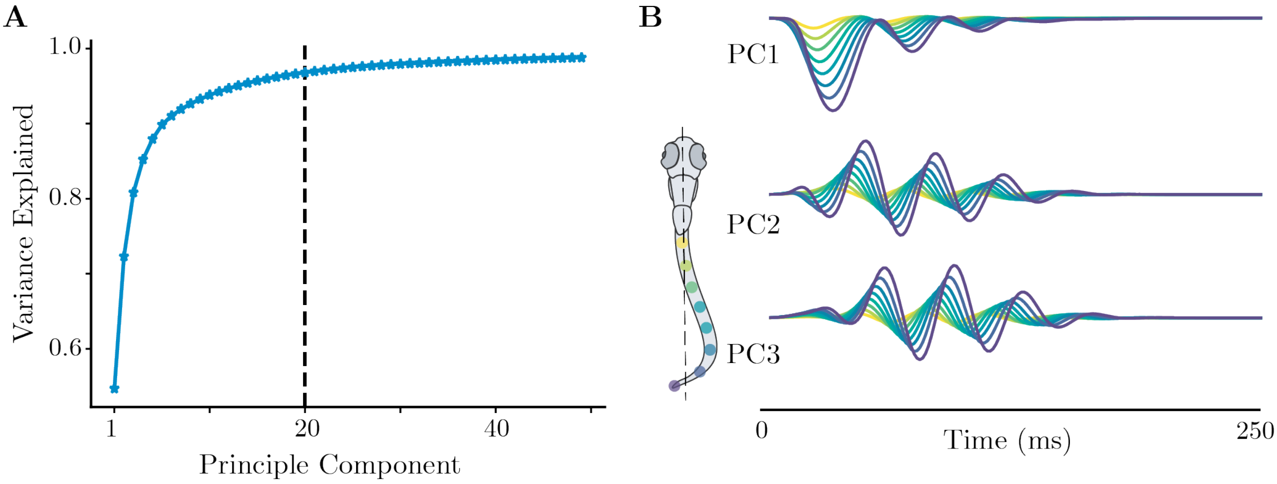


**Figure S1. Principle component analysis (PCA) of bout space. (A)** All bouts are represented by the cumulative tail angle of 8 points on the tail for $250$ms from the start of the bout. For the dataset from Marques et al. (2018), a single bout is then composed of $175$ frames $\times$ $8$ (at a sampling frame rate of 700Hz). We randomly sample 25 recordings across all sensory conditions ($\approx 50,000$ bouts per sampling), estimate a covariance matrix and obtain its eigenvalues and eigenvectors. We then estimate the mean covariance matrix and the means of the eigenvalues and eigenvectors across all such possible samples of the data. We find that the first 20 eigenvectors capture $>$95% of the variance. We also calculate 95% errorbars on the estimate of the variance explained across multiple resampling, but these are too minuscule to notice. **(B)** To showcase what the PCA eigenvectors represent, we plot the first three principle components. The first component PC1 mostly captures turning biases (leftward vs rightward turns), the second and third components, PC2 and PC3, correspond to forward waves. Final scale details of rarer bouts are captured by the remaining modes.

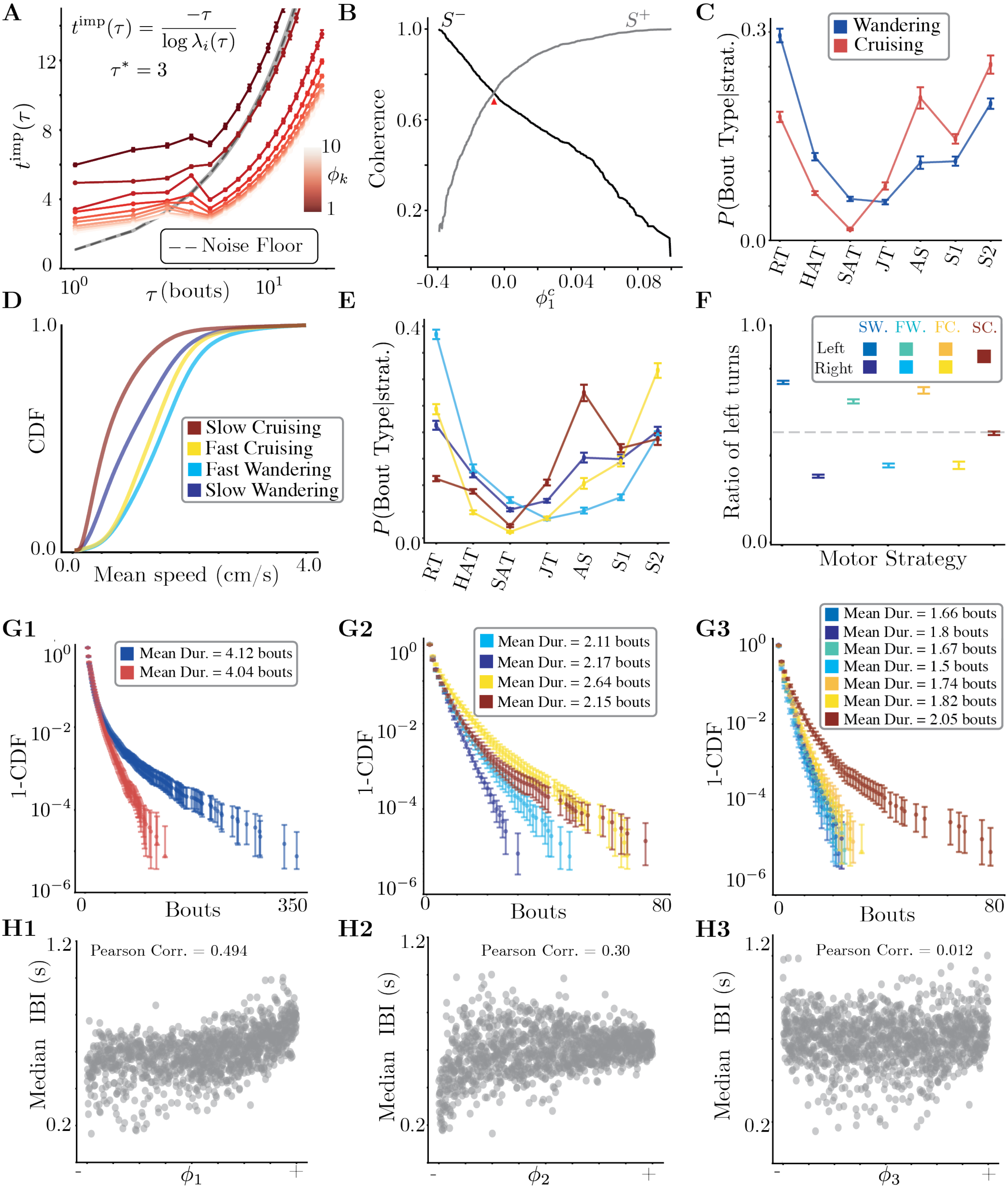
A
$t^{\mathrm{imp}}(\tau) = \frac{-\tau}{\log \lambda_i(\tau)}$
$\tau^* = 3$
$t^{\mathrm{imp}}(\tau)$
$\tau$ (bouts)
$\phi_k$
Noise Floor
B
Coherence
$S^-$
$S^+$
$\phi_1^c$
C
$P$(Bout Type|strat.)
Wandering
Cruising
RT HAT SAT JT AS S1 S2
D
CDF
Mean speed (cm/s)
Slow Cruising
Fast Cruising
Fast Wandering
Slow Wandering
E
$P$(Bout Type|strat.)
RT HAT SAT JT AS S1 S2
F
Ratio of left turns
Motor Strategy
SW. FW. FC. SC.
Left
Right
G1
1-CDF
Bouts
Mean Dur. = 4.12 bouts
Mean Dur. = 4.04 bouts
G2
1-CDF
Bouts
Mean Dur. = 2.11 bouts
Mean Dur. = 2.17 bouts
Mean Dur. = 2.64 bouts
Mean Dur. = 2.15 bouts
G3
1-CDF
Bouts
Mean Dur. = 1.66 bouts
Mean Dur. = 1.8 bouts
Mean Dur. = 1.67 bouts
Mean Dur. = 1.5 bouts
Mean Dur. = 1.74 bouts
Mean Dur. = 1.82 bouts
Mean Dur. = 2.05 bouts
H1
Median IBI (s)
Pearson Corr. = 0.494
$\phi_1$
H2
Median IBI (s)
Pearson Corr. = 0.30
$\phi_2$
H3
Median IBI (s)
Pearson Corr. = 0.012
$\phi_3$

**Figure S2. Details of extracting and analyzing the long lived modes from the transition matrix $T_{\text{ensemble}}$ (A)** Implied timescales $t_{\text{imp}}(\tau)$ as a function of the transition time $\tau$. We estimate the implied timescales by randomly sampling 7500 bouts from each of the 14 conditions over 100 random seeds. Errorbars are bootstrapped 95% estimates of the implied timescales. For each value of $\tau$, we estimate a transition matrix by counting the number of times the dynamics goes from state $s_i$ to $s_j$ after $\tau$ bouts. When $\tau$ is too large, the dynamics starts to effectively mix, meaning that samples that are too disparate in time are practically independent from each other. In this limit, there should only be one surviving non-zero eigenvalue of $T_{ij}$, corresponding to the steady-state distribution. However, the finite length of the recordings induces a non-zero noise floor, which we gauge by estimating the largest eigenvalue from a transition matrix built from shuffled symbolic sequences (gray line). At smaller $\tau$, a large fraction of eigenvalues are significant, and we choose an intermediate $\tau^* = 3$ bouts to isolate the slow modes from the bulk of the spectrum. **(B)** Coherence of each metastable strategy a function of $\phi_1^c$. We scan along $\phi_1$ and define $S^-$ and $S^+$ as collections of microstates that have $\phi_1$ smaller or larger than a threshold $\phi_1^c$, respectively. We then estimate the fraction of probability that remains within both sets as we vary $\phi_1^c$, which we call coherence (see Methods for details) Froyland (2005). We identify the longest lived motor strategies by identifying the value of $\phi_1^c$ that maximizes the coherence of both motor strategy. **(C)** Probability of observing different bout types in each of the coarse-grained strategies: Cruising and Wandering. We find a bias towards turning behaviors in Wandering (**R**outine **T**urn, **H**igh **A**ngle **T**urn, **S**pot **A**voidance **T**urn), while Cruising behaviors are dominated by forward movements (**A**pproach **S**wim, **S**low **1**, **S**low **2**). Notice also that despite these broad differences, there is also a non-negligible fraction of routine turns (RT) during Cruising, as well as a large fraction of S2 in Wandering. **(D)** Cumulative Distribution Function (CDF) of the mean instantaneous speed in a bout sequence in the four long-lived motor strategies. We discover finer scale states by using faster decaying eigenvectors: here, we use $\phi_1$ and $\phi_2$ to reveal 4 motor strategies, which correspond to Slow and Fast variations of Cruising and Wandering (see Methods for details). **(E)** Probability of observing different bout types in each of the 4 motor strategies. **F** We further identify faster timescale motor strategies using the first three eigenvectors $\phi_k, k \in 1, 2, 3$ (see Methods for details). We find left and right direction bias variations of the slow wandering, fast wandering and fast cruising states. In contrast, the slow cruising does not exhibit persistent left/right biases. **(G1)** Complementary cumulative distribution function (1 - CDF) of the time spent in the Cruising and Wandering states across all fish. We find a broad distribution of timescales, with a comparatively small average sequence length of $4.12$ bouts $\Rightarrow 3.17\,\text{s} \pm (0.11, 20.59)$ s in Wandering and $4.04$ bouts $\Rightarrow 2.77\,\text{s} \pm (0.11, 17.12)$ s in Cruising. Errorbars represent bootstrapped 95% confidence intervals across fish. **(G2)** Complementary cumulative distribution function (CCDF) of the time spent in the slow and fast variation of Cruising and Wandering states across all fish. As expected, these states capture faster timescale properties of the bout sequence dynamics, with slow cruising lasting on average $1.03\,\text{s} \pm (0.11, 5.16)$ s, fast cruising $1.35\,\text{s} \pm (0.11, 9.61)$ s, slow wandering $1.32\,\text{s} \pm (0.11, 7.33)$ s and fast wandering $1.35\,\text{s} \pm (0.11, 9.12)$ s. Errorbars represent 95% confidence intervals measure across fish. **(G3)** Complementary cumulative distribution function (CCDF) of the time spent in the left/right variation of the Slow/Fast strategies. Errorbars represent 95% confidence intervals measure across fish. **(H1)** Median inter-bout interval in the bout sequences corresponding to a given microstate, plotted against $\phi_1$. We find that the inter-bout interval is correlated with $\phi_1$. **(H2)** Median inter-bout interval in the bout sequences corresponding to a given microstate, plotted against $\phi_2$. We find that slow microstates have lower inter-bout intervals. **(H3)** Median inter-bout interval in a microstate plotted against $\phi_3$. There is almost no correlation between the inter-bout interval and the egocentric direction preference.

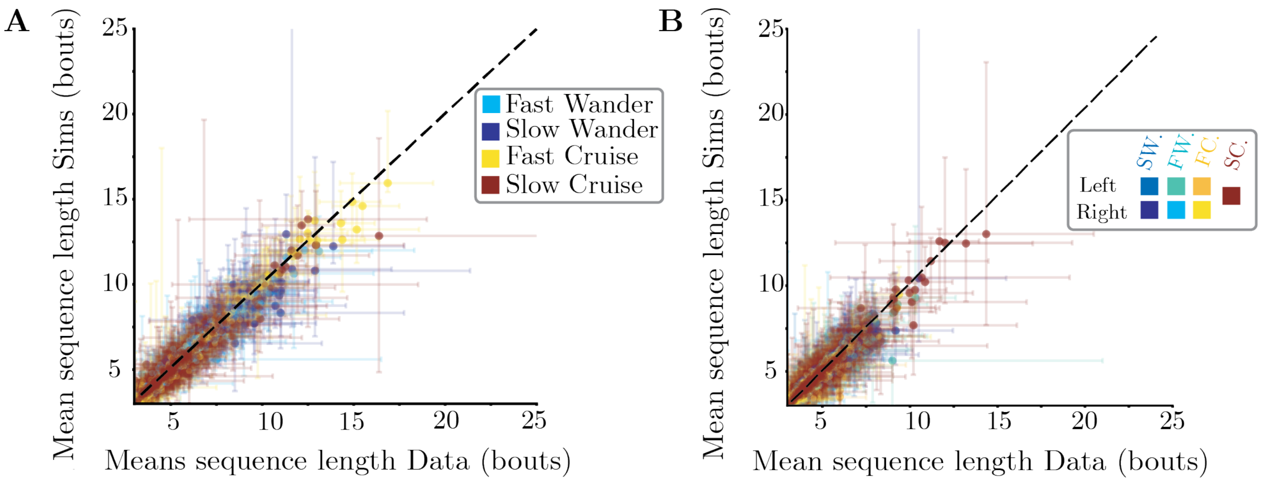


**Figure S3. Simulations for individual fish at different coarse-graining scales (A)** Mean sequence length for data and simulations at the level of four metastable states. **(B)** Mean sequence length for data and simulations at the level of seven metastable states. We get a good match between the data and the simulations even at the scale of these shorter timescale strategies. Errorbars represent bootstrapped 95% confidence intervals on the mean sequence length. To obtain error bars from the simulations, we collect behavioral events from 100 independent simulations, and bootstrap by sampling a number of events equal to the one observed in the real data (see Methods).

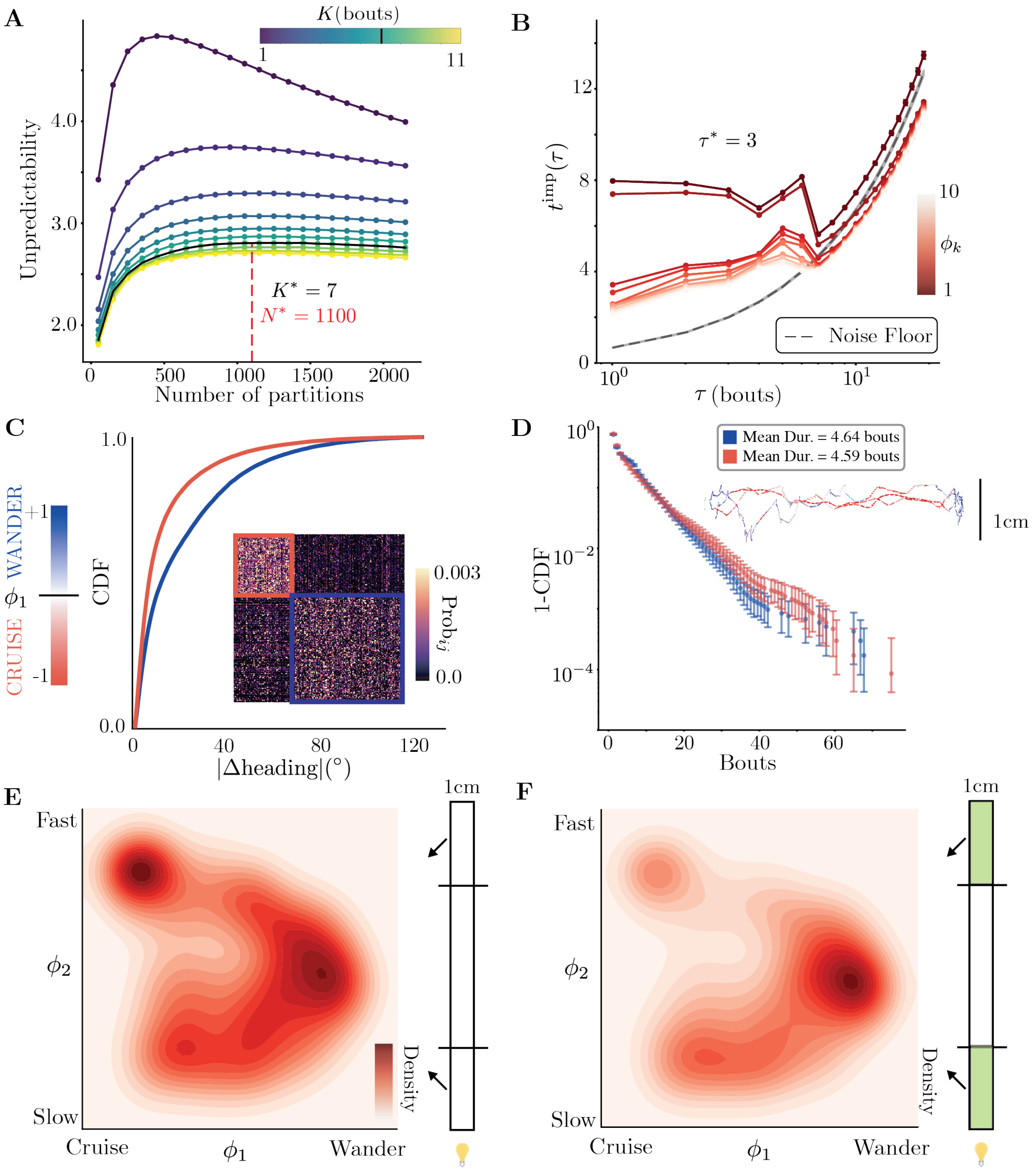
A
K (bouts)
1
11
Unpredictability
4.0
3.0
2.0
K* = 7
N* = 1100
0
500
1000
1500
2000
Number of partitions
B
12
8
4
0
τ* = 3
t^imp(τ)
10
ϕ_k
1
Noise Floor
10^0
10^1
τ (bouts)
C
1.0
0.0
CDF
CRUISE ϕ_1 WANDER
+1
-1
0.003
Prob_ij
0.0
0
40
80
120
|Δheading|(°)
D
10^0
10^-2
10^-4
1-CDF
Mean Dur. = 4.64 bouts
Mean Dur. = 4.59 bouts
1cm
0
20
40
60
Bouts
E
Fast
ϕ_2
Slow
Cruise
ϕ_1
Wander
Density
1cm
F
Fast
ϕ_2
Slow
Cruise
ϕ_1
Wander
Density
1cm

**Figure S4. Discovering the long lived modes of behavior in the dataset from Reddy et al. (2022).** We note that this dataset was collected at a lab different from the one that collected the data used in the main text Marques et al. (2018), and the bouts were tracked using a different tracking algorithm (Zebrazoom Mirat et al. (2013)). Here, the fish are swimming in a $1\,\mathrm{cm} \times 14\,\mathrm{cm}$ arena or were subject to aversive acidic pH. We take all recordings where the fish swim for longer than 250 bouts, which gives 123 recordings without pH and 93 recordings with pH (see Methods). **(A)** Unpredictability as a function of the number of stacked bouts $K$ and the number of partitions. We measure unpredictability as in the main text: we equally sample 40,000 bouts from the two different sensory condition (with and without acidic pH), and estimate the entropy rate of the Markov chain constructed by partitioning the space defined by sequences of $K$ bouts into $N$ microstates. We find that in this dataset $K^* = 7$ bouts minimizes the entropy rate, and $N^* = 1100$ is the maximum number of partitions beyond which finite-size effects results in an underestimation of the entropy rate. Notice that, compared to Fig. 1, we find a larger $K^*$, reflecting longer timescale correlations among the tracked bouts. Errorbars represent bootstrapped 95% confidence intervals over 50 resamples of the data. **(B)** Implied timescales of the transition matrix at $K^* = 7, N^* = 1100$. We choose $\tau^* = 3$ as before. 40,000 microstates are sampled from each condition. Errorbars represent bootstrapped 95% confidence intervals over 20 resamplings of the symbolic sequences. **(C)** As in the main results, the first non-trivial eigenvector $\phi_1$ of the ensemble transition matrix coarse-grains the dynamics into Cruising-Wandering strategies. We plot the cumulative distribution of the mean absolute change in heading in each motor strategy. (inset) The coarse-graining results in a block-diagonal structure in the transition matrix. **(D)** Complementary cumulative distribution functions (1-CDF) of the sequence length in either Cruising or Wandering strategies with one example trajectory of 200 bouts color coded by $\phi_1$. **(E)** Kernel density estimates along $\phi_1, \phi_2$ for the non-acidic control recordings, sampled only in regions where pH would have spread. Fish have a distributed preference for both cruising and wandering in these regions in the recordings. **(F)** Kernel density estimates along $\phi_1, \phi_2$ for the acidic pH recordings, sampled only in regions where pH would have spread. Fish have a specific preference for wandering in these regions, pointing to wandering as a long lived response to the pH.

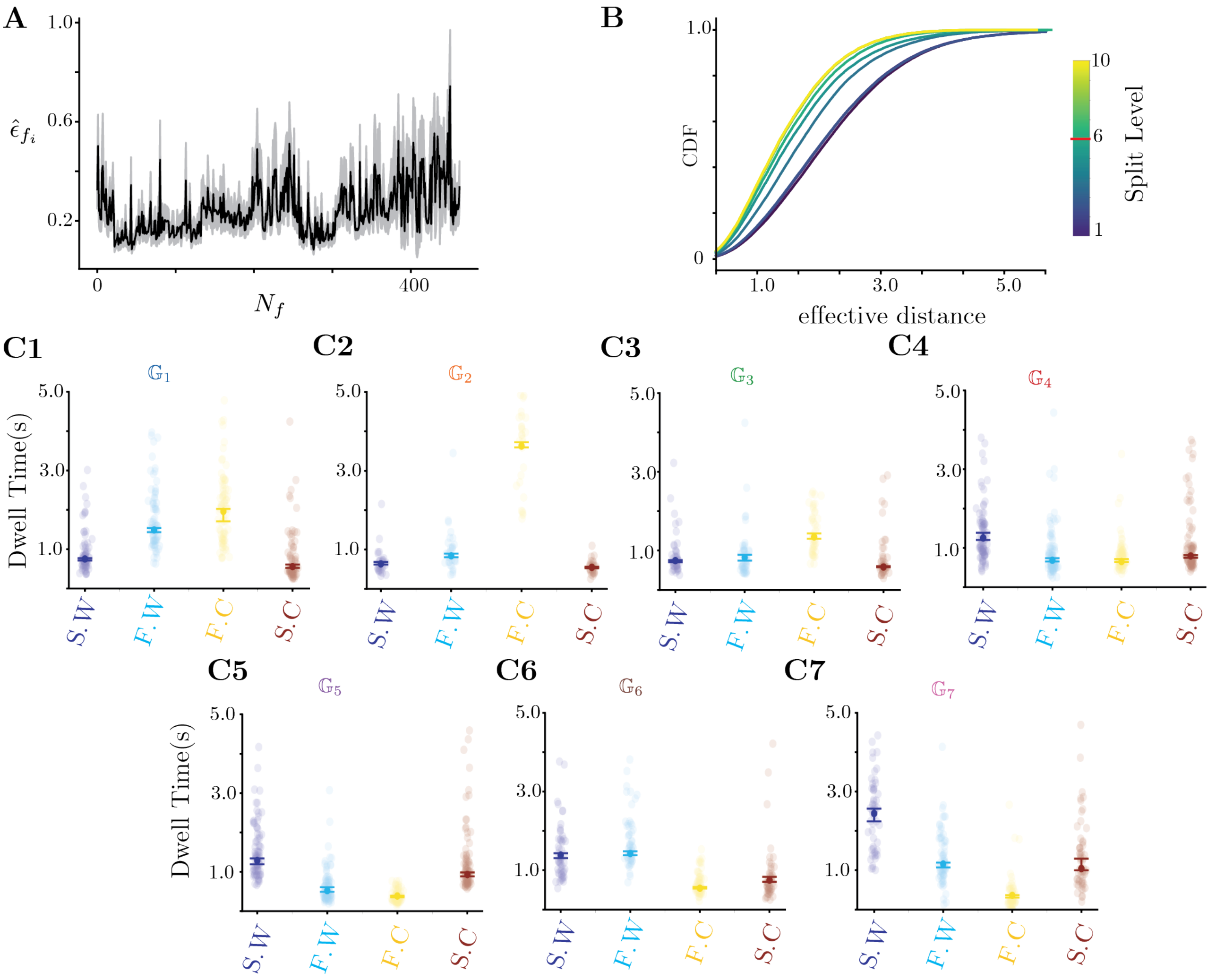


**Figure S5. Details on the Hierarchical multiplicative diffusive clustering and the discovered phenotypic groups** $\mathbb{G}$ **(A)** The effective scale separation $\hat{\epsilon}_{f_i}$ for each fish at $q = 7$. Light grey represent 95% bootstrapped confidence intervals. **(B)** Cumulative distributions of effective distances within a cluster at different iterations $L$ of the hierarchical clustering. The distributions change very little after $L = 6$, setting that as the cut-off point for the clustering. **(C)** Each panel shows the median dwell time across fish in either fast wandering, slow wandering, fast cruising or slow cruising for one of the four behavioral groups $\mathbb{G}$. We calculate the mean in each metastable strategy within a fish and take medians across fish. Each point represents the mean dwell time of a single fish, errorbars represent bootstrapped 95%confidence intervals on the median across fish. **(C1)** Median dwell times in $\mathbb{G}_1$. Slow Wandering: $0.75\,(0.71, 0.77)$ s, Fast Wandering: $1.49\,(1.44, 1.59)$ s, Fast Cruising: $1.96\,(1.71, 2.03)$ s, Slow Cruising: $0.56\,(0.53, 0.61)$ s **(C2)** Median dwell times in $\mathbb{G}_2$. Slow Wandering: $0.65\,(0.62, 0.68)$ s, Fast Wandering: $0.84\,(0.8, 0.89)$ s, Fast Cruising: $3.63\,(3.59, 3.72)$ s, Slow Cruising: $0.55\,(0.52, 0.56)$ s **(C3)** Median dwell times in $\mathbb{G}_3$. Slow Wandering: $0.74\,(0.71, 0.76)$ s, Fast Wandering: $0.82\,(0.74, 0.89)$ s, Fast Cruising: $1.35\,(1.3, 1.44)$ s, Slow Cruising: $0.58\,(0.57, 0.6)$ s **(C4)** Median dwell times in $\mathbb{G}_4$. Slow Wandering: $1.25\,(1.2, 1.38)$ s, Fast Wandering: $0.69\,(0.65, 0.74)$ s, Fast Cruising: $0.65\,(0.64, 0.71)$ s, Slow Cruising: $0.8\,(0.75, 0.82)$ s **(C5)** Median dwell times in $\mathbb{G}_5$. Slow Wandering: $1.3\,(1.19, 1.34)$ s, Fast Wandering: $0.52\,(0.49, 0.61)$ s, Fast Cruising: $0.39\,(0.35, 0.39)$ s, Slow Cruising: $0.93\,(0.87, 0.99)$ s **(C6)** Median dwell times in $\mathbb{G}_6$. Slow Wandering: $1.4\,(1.3, 1.43)$ s, Fast Wandering: $1.42\,(1.38, 1.48)$ s, Fast Cruising: $0.54\,(0.53, 0.57)$ s, Slow Cruising: $0.75\,(0.71, 0.85)$ s **(C7)** Median dwell times in $\mathbb{G}_7$. Slow Wandering: $2.44\,(2.19, 2.57)$ s, Fast Wandering: $1.16\,(1.07, 1.19)$ s, Fast Cruising: $0.36\,(0.31, 0.37)$ s, Slow Cruising: $1.04\,(1.0, 1.29)$ s

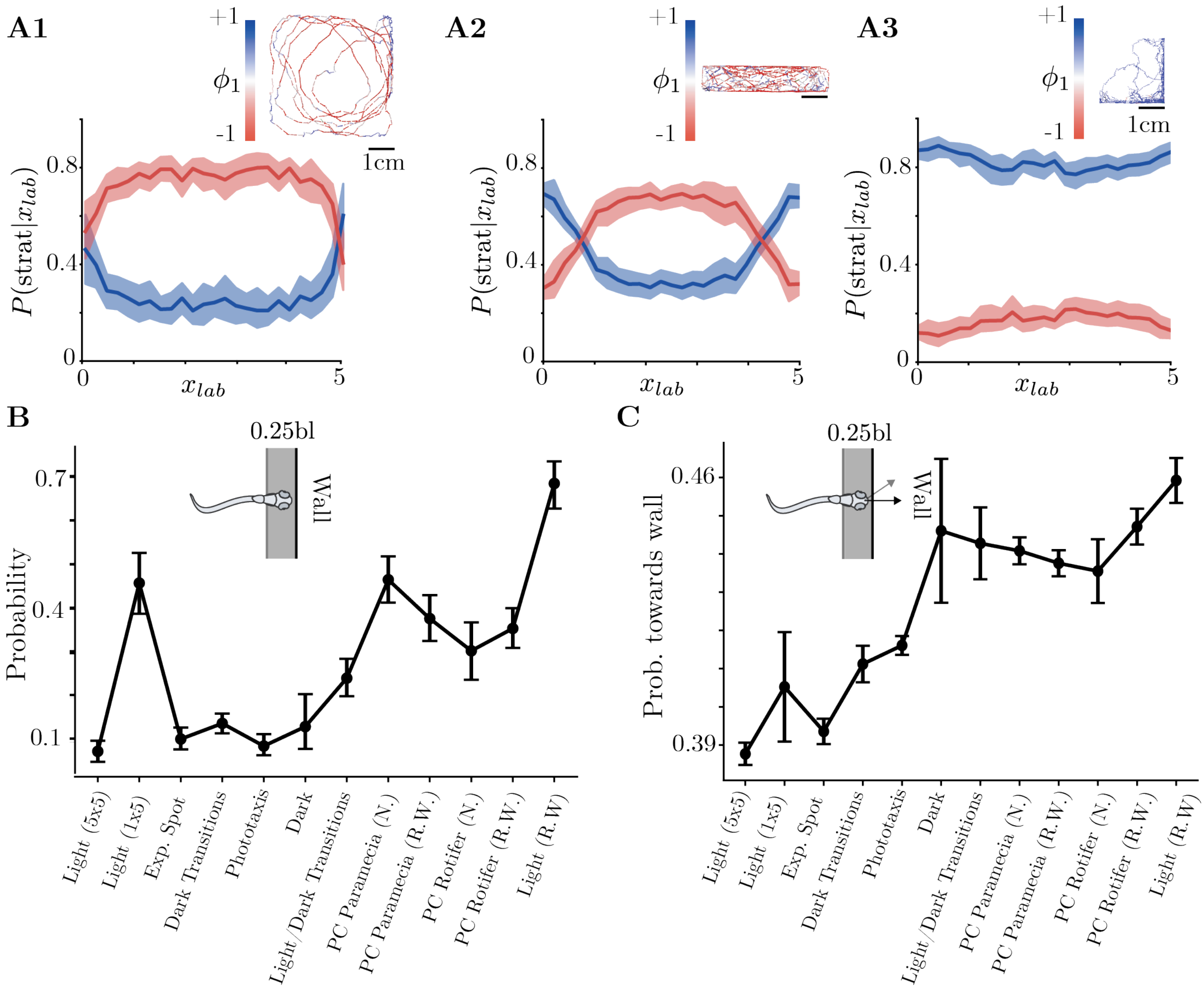


**Figure S6. Spatial nature of the usage of Cruising-Wandering states, and details of the wall-following behavior across free-swimming conditions (A)** Probability of executing either Cruising or Wandering strategies conditioned on the position along the horizontal axis $x_{\mathrm{lab}}$. As an inset, we plot an example trajectory of 500 bouts in each condition, color coded by $\phi_1$. **(A1)** In a 5cm x 5cm arena, Wandering prevails only when the fish is at the corners of the arena, where it is forced to engage in a sequence of reorientations. **(A2)** In a 1cm x 5cm arena, similar to the 5cm x 5cm arena, the probability of Wandering increases only when the fish is at the corners. Shortening the $y-$axis of the arena prevents the fish from cruising along the vertical axis, whereas in the 5cm x 5cm arena fish could still cruise even for small and large values of $x_{\mathrm{lab}}$. **(a3)** In a 2.5cm x 2.5cm arena, fish that were raised with rotifers engage in Wandering behavior throughout the entire arena. Notice also how in this case fish barely explore the center of the arena, mostly interacting with the walls while Wandering. **(B)** Probability of thigmotaxis in each condition (here defined as the proportion of bouts in which the head position is on average smaller that a quarter of a body length away from the wall). Fish that were raised with Rotifers but are freely swimming have the highest probability of thigmotaxis, followed by the remaining prey capture conditions. In addition, we notice that the Light (1x5) condition the fish also spend time close to the wall, but they do so mostly by cruising along the major axis of the arena, see panel A2. **(C)** Probability of the reorienting towards the wall when the fish is close to the wall. Unlike the other Light conditions, fish raised with prey have a much higher probability of orienting towards the wall during the behavior, indicating a different role to their thigmotatic behavior. Additionally, we find that in other prey capture conditions, as well as in the Dark and Ligh/Dark Tx. condition (in which Wandering strategies are also prevalent), fish also reorient towards the wall when they are close to it.

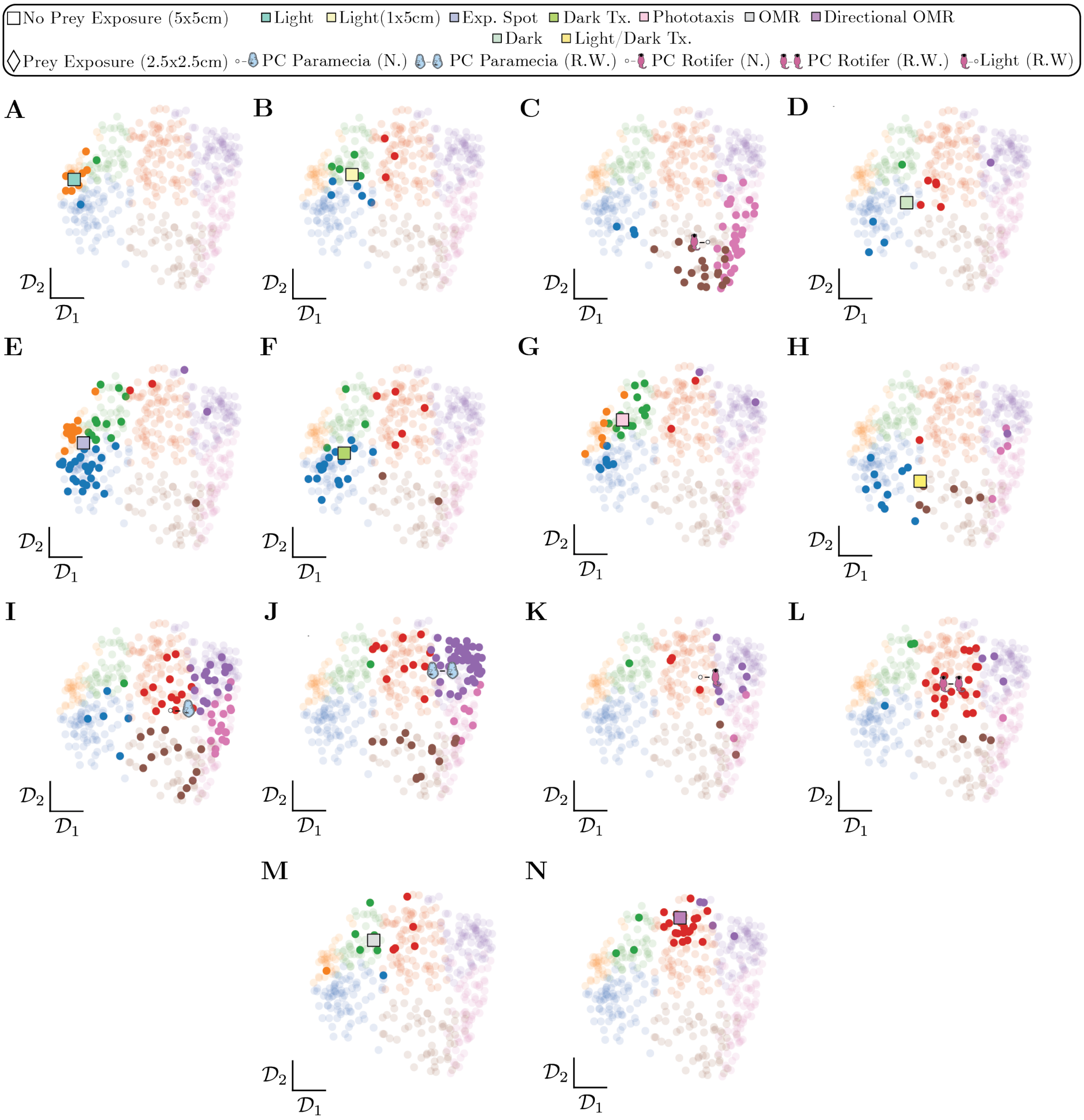


**Figure S7. Variability among fish from different sensory conditions.** In each panel, we plot the position of each individual fish along $\mathcal{D}_1$ and $\mathcal{D}_2$ of the transition matrix space as the background, color coding each fish by their respective behavioral group. In addition, we highlight in darker colors the fish corresponding to each sensory condition, as well as the transition matrix obtained from all fish in a given condition through a different tick mark. **(A)** Light (5cm x 5cm) condition. **(B)** Variability among fish in the Light (1cm x 5cm) condition **(C)** Variability among fish in the Light R.W (2.5cm x 2.5cm) condition. **(D)** Dark (5cm x 5cm) condition. **(E)** Expanding Spot (5cm x 5cm) condition. **(F)** Dark transitions (5cm x 5cm) condition. **(G)** Phototaxis (5cm x 5cm) condition. **(H)** Light/Dark transitions (5cm x 5cm) condition. **(I)** Prey Capture Paramecia Naive (2.5cm x 2.5cm) condition. **(J)** Prey Capture Paramecia Raised With (2.5cm x 2.5cm) condition. **(K)** Prey Capture Rotifer Naive (2.5cm x 2.5cm) condition. **(L)** Prey Capture Paramecia Rotifer Raised With (2.5cm x 2.5cm) condition. **(M)** OMR (1cm x 5cm) condition. **(N)** Directional OMR (5cm x 5cm) condition.

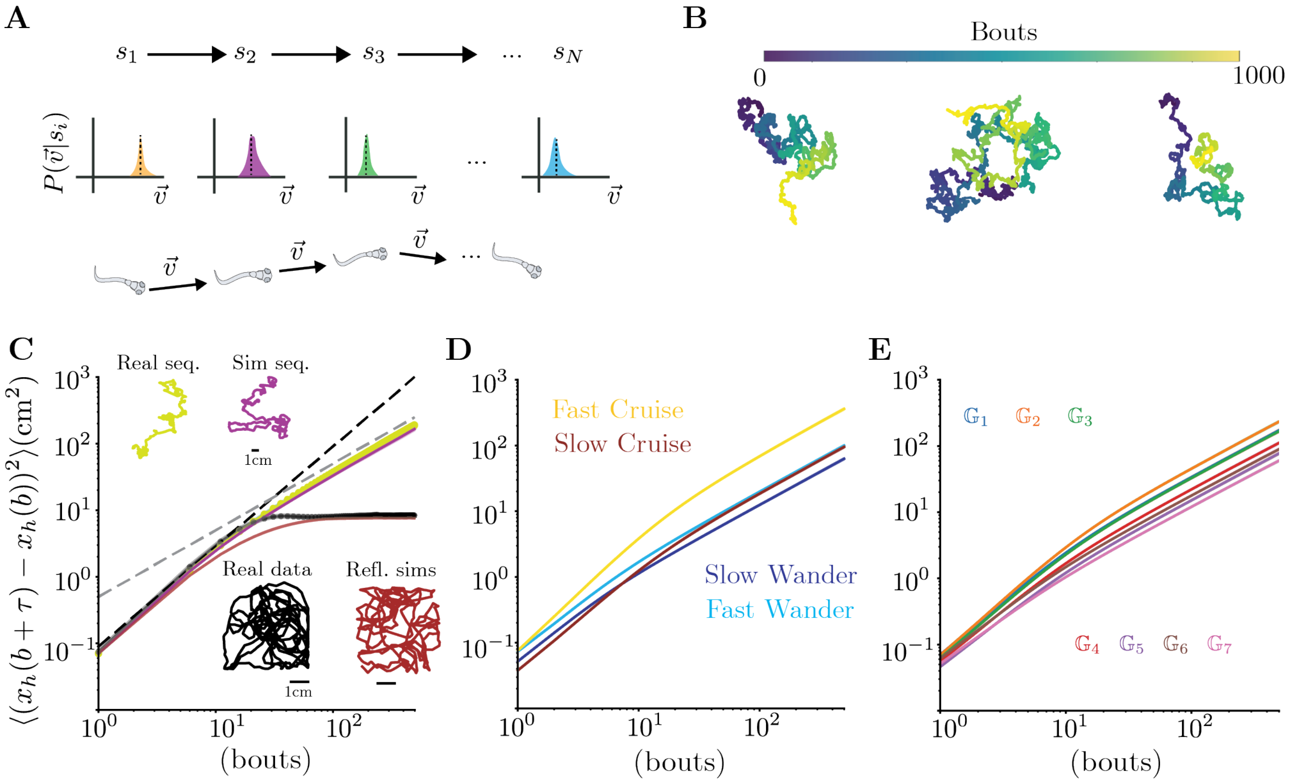


**Figure S8. Assessing the statistical properties of the simulated trajectories through the mean squared displacement. (A)** Schematic of the simulation procedure. From a transition matrix among all $N^* = 1200$ states and a discrete symbol $s_i$, we sample the next symbol $s_j$ from the $i$-th row of transition matrix, which corresponds to the conditional probability of observing any state $s_j$ given state $s_i$. For each symbol $s_i(t)$, we then sample a velocity vector from the distribution of velocities observed when fish where in state $s_i$ (see Methods for details). **(B)** Three example trajectories from the simulation procedure, each 1000 bouts long. **(C)** We estimated the mean squared displacement (MSD) from the data (shown as a black scatter plot), as well as different kinds of trajectory simulations (colored lines). In the data, we observe super-diffusive behavior on short time scales, with the MSD scaling approximately as MSD $\sim \tau^{1.5}$ (black dashed line). However, the finite-size of the arenas quickly induces strong finite-size effects which bound the MSD from above. As a first assessment of the effects of the arena wall, we tried to reconstruct fish trajectories without boundary constraints by using the real symbolic sequence of each fish to generate new trajectories. We sample velocity vectors from the distribution of velocities obtained from the bouts corresponding to each symbol and use them to generate artificial trajectories, which we label as Real seq. (yellow), see schematic of Fig. S8A and Methods for details. Notably, such trajectories match the statistics of the real data up to $\approx 20$ bouts, at which point the data reaches its upper bound while the Real seq. trajectories start entering a diffusive regime in which MSD $\sim \tau$ (gray dashed line). We then generate trajectories from simulated symbolic sequences, which we call Sim seq. (purple). We find that the trajectories obtained from simulated symbolic sequences closely match the data on short timescales, but also match the trajectories obtained from the real symbolic sequence well across scales. Overall, we find that the arena sizes severely challenges an accurate estimate of the diffusive properties of larval zebrafish navigation, but that nonetheless our simulations match the statistics observed in the data within its limits. Note also that simply enforcing reflective boundary conditions (red) is not enough to capture the statistics, even at short timescales, pointing to an active interaction of the fish with the wall. **(D)** Mean squared displacements of simulations from $N = 4$ metastable strategies. **(E)** Mean squared displacements of simulations from behavioral groups $\mathbb{G}_i, i \in [1, 7]$.
